# Structures of the germline-specific Deadhead and Thioredoxin T proteins from *Drosophila melanogaster* reveal unique features among Thioredoxins

**DOI:** 10.1101/2020.07.29.226944

**Authors:** Regina Freier, Eric Aragón, Błażej Bagiński, Radoslaw Pluta, Pau Martin-Malpartida, Lidia Ruiz, Miriam Condeminas, Cayetano Gonzalez, Maria J. Macias

## Abstract

Thioredoxins (Trxs) are ubiquitous enzymes that regulate the redox state in cells. In *Drosophila*, there are two germline-specific Trxs, Deadhead (Dhd) and TrxT. Both proteins belong to the L(3)mbt malignant brain tumor signature and to the MMS survival network of genes that mediate the cellular response to DNA damage. Dhd is a maternal protein required for early embryogenesis that promotes protamine-histone exchange in fertilized eggs and midblastula transition. TrxT is testis-specific and associates with the lampbrush loops of the Y chromosome.

Here we present the first structures of Dhd and TrxT that unveil new features of these Thioredoxins. Dhd is highly positively charged, unusual in canonical Trxs. This positively charged surface can facilitate its approximation to DNA and to protamine oligomers, to promote chromatin remodeling. On the other hand, TrxT contains a C-terminal extension, mostly unstructured and highly flexible, which wraps the conserved core through a closed conformation. This extension partially covers the catalytic site and modulates the redox activity of the protein.

The information provided by these structures can guide future work aimed at understanding how redox inputs modulate the initial steps of embryo development in *Drosophila* and may help in the design of molecular inhibitors through a structure-based approach.

**Highlights:**

1. We have determined the first structures of the germline-specific Trxs Dhd and TrxT.
2. Dhd has a highly positively charged surface that facilitates its approximation to DNA and protamine oligomers, to promote chromatin remodeling.
3. TrxT contains a C-terminal extension, highly unusual in canonical Trxs, mostly unstructured and highly flexible.
4. The TrxT C-terminal extension partially covers the catalytic site and modulates the redox activity of the protein.
5. The differences observed in Thioredoxins can help in fine-tuning specific molecules to be active against selected insect species.

## INTRODUCTION

Thioredoxins (Trxs) are present in all living organisms and cellular compartments and they form the most numerous subfamily of oxidoreductase enzymes in nature (1). In addition to their general role in controlling redox homeostasis in cells, Trxs participate in specific tasks, including the regulation of programmed cell death and transcription factor activity and the modulation of inflammatory responses, and they also serve as growth factors (2). Trx domains also contribute to protein folding and prevent protein aggregation. An example of this role is represented by the Protein Disulfide-Isomerase family (PDI), which regulates protein misfolding by catalyzing the formation and breakage of disulfide bonds during proteins synthesis (3).

Trxs share a conserved fold consisting of five beta-strands and four alpha-helices and a catalytic motif with two conserved Cys residues. The catalytic motif (Cys-X-X-Cys) is located at the beginning of the second helix, on the protein surface, and it is partially exposed to facilitate access to substrates (Trx-2, Supplementary Figure 1A,B) (4). The redox mechanism modulated by Trxs is a coordinated reaction where a substrate protein and the Trx/TrxR system act in an orchestrated manner (schematically represented in Figure 1A). The redox cycle starts with a reduced form of Trx and the catalytic Cys (Cys32) in the form of a thiolate (Supplementary Figure 1A). This state is stabilized by a hydrogen bond to the second Cys of the motif (Cys35-SH group). The thiolate is then able to form a transient inter-molecular disulfide bond with the Cys present in the oxidized substrate. The catalytic cycle ends when Cys35 in the enzyme attacks this inter-molecular disulfide and forms a new intra-molecular bond with Cys32, releasing the reduced substrate and the oxidized enzyme (5). Trx recovers its initial state by the action of TrxR, which reduces Trx using NADPH/FAD as the source of reducing equivalents (Figure 1A) (2). Furthermore, Trx and Trx Reductase (TrxR) are overexpressed in many tumor cells whose proliferation is dependent on a high supply of deoxyribonucleotide (6,7). Hence, inhibition of the Trx/TrxR system emerged as an attractive target for anticancer drugs to induce cell death (8). In this regard, the use of *Drosophila* as a model system for high-throughput screening of molecules has attracted the attention of researchers (9).

**Figure 1.**
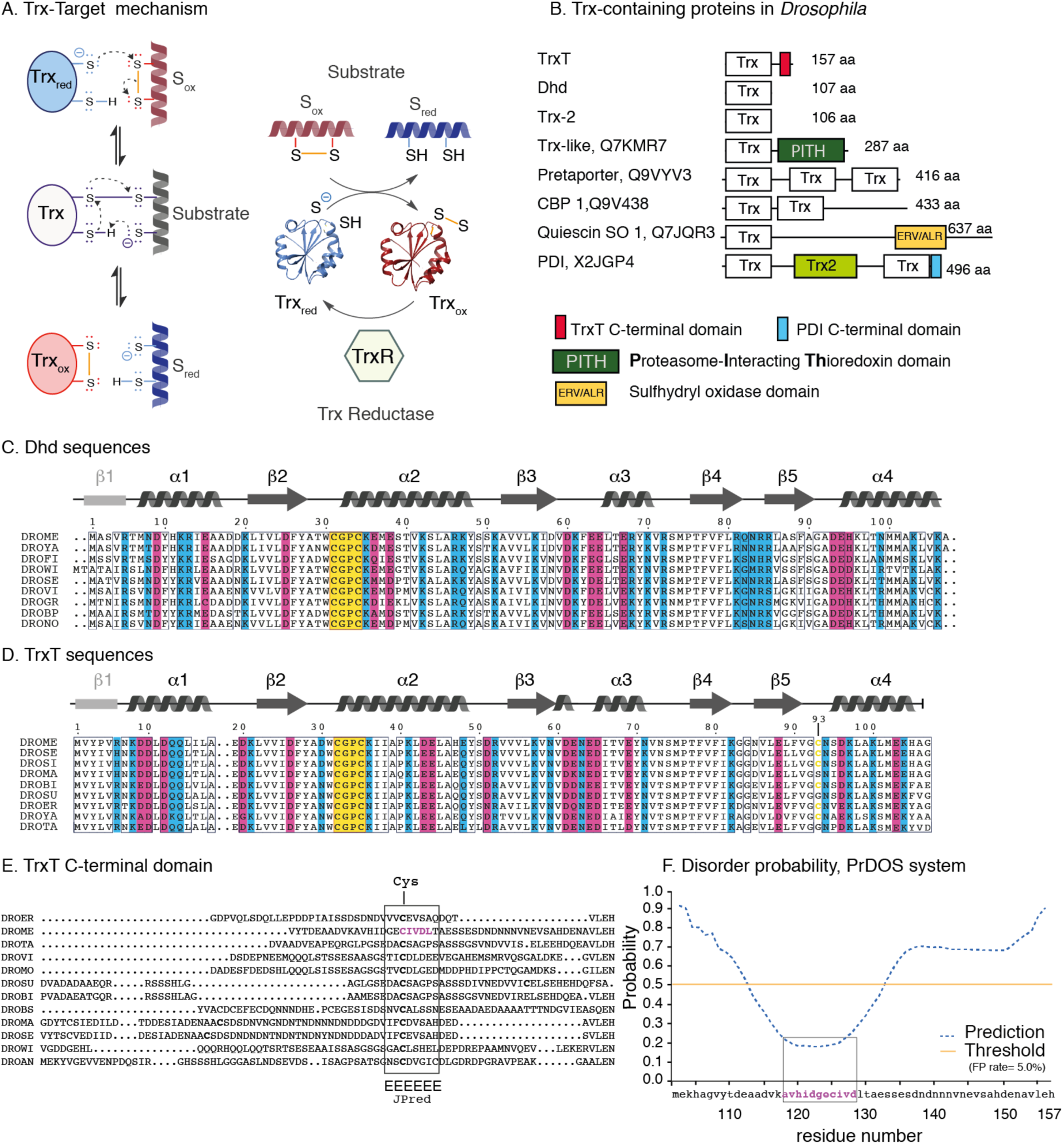
Redox mechanism and Trx proteins in *D. melanogaster*. **A.** Schematic description of the Trx-TrxR redox mechanism adapted from (2). **B.** Trx-containing proteins in *Drosophila melanogaster.* UniProt codes and domains are indicated. Abbreviations: CBP: calcium-binding protein, PDI: protein disulfide isomerase, SO: sulfhydryl oxidase. Isoelectric Points for the different Trx domains are as follows: (6.5, Q7KMR7), (5.0, Q9VYV3, three domains), (6.6, Q9V438, two domains), (4.9, Q7JQR3) and (5.1, X2JGP4, two domains). A sequence alignment of these domains is shown as Supplementary Figure 1B. **C.** Alignment of selected Dhd protein sequences. An extended version of this alignment is depicted in Supplementary Figure 1D. The species are named with acronyms. RefSeq codes and species are indicated in Table 2. The catalytic region is indicated with a yellow box and conserved positively and negatively charged residues are highlighted as blue and purple bars, respectively. Secondary structure elements based on the *Drosophila melanogaster* structure determined in this work are shown on top of the alignment. To facilitate comparison to other Trx structures, the β1 strand is indicated in gray. **D.** Alignment of selected TrxT proteins. An extended version of this alignment is depicted in Supplementary Figure 1E. Names and RefSeq codes are shown in Table 3. Color patterns as in C. **E.** Sequence comparison of the TrxT C-terminal domain. Boxed region indicates the cysteine-containing motif predicted to adopt an extended conformation by the software JPred (28). The sequence that forms a disulfide bond with Cys93 and adopts an extended conformation in the crystals is indicated in pink. **F.** Disorder probability calculated for the C-terminal domain using the PrDOS server (http://prdos.hgc.jp). Residues highlighted in pink are predicted to adopt ordered conformations.

In *Drosophila melanogaster*, there are several Trxs. Trx-2 (also known as Dm Trx) is a non-essential protein, similar to human Trx1, and widely distributed in all cellular compartments. On the contrary, the female germline-specific Deadhead (Dhd) protein and the male germline-specific TrxT have highly specific distribution and functional roles (Figure 1B) (10). Both TrxT and Dhd belong to the *l(3)mbt* malignant brain tumor signature genes (*lethal(3)malignant brain tumor*) (11,12) and they have been identified as part of the ‘survival network’ of genes that mediate the cellular response to DNA damage induced by alkylating agent methyl methanesulfonate (MMS) (13). TrxT has exclusive functions, as its association with the Y chromosome lampbrush loops (10). Dhd function is essential for *Drosophila* embryo development and *Dhd*^-^ mutant oocytes show meiotic defects (14). Dhd is also required for early embryogenesis and metabolic remodeling, and it participates in the redox control of protamines and in sperm chromatin remodeling *in vivo* (15,16). All these roles are exclusive to Dhd, since the ubiquitous Trx2 cannot recognize these substrates (17). Moreover, Dhd plays crucial roles in the oocyte-to-embryo transition, where it reduces and modulates the activity of ribosomal and RNA-binding proteins, as well as that of the histone demethylase NO66 protein (18). Recently, it has also been reported that the transcriptional regulation of Dhd is modulated by the lysine-specific demethylase 5 (KDM5), a potent chromatin remodeler during female gametogenesis (19).

From the sequence perspective, we noted that both Dhd and TrxT have key features that distinguish them from other Trxs. Dhd has an unusually high number of Arg/Lys residues in its sequence whereas TrxT has a C-terminal domain of approximately 50 residues with yet unknown fold and function (Supplementary Figure 1C) (20). These two characteristics suggest that each protein has exceptional structural properties that correlate with their specific functional roles. To fill this knowledge gap regarding the structures of these germline-specific Trxs and to expand the potential application of *Drosophila* as a model organism for studying redox regulation, we turned our attention to the structures of the nuclear Trxs TrxT and Dhd. Using X-ray crystallography, we found that both proteins have a conserved fold and catalytic site but that they display structural properties that correlate with their specific sequences, thereby illustrating the versatility of the conserved Trx fold to fine-tune its function.

Our data revealed that Dhd has positively charged patches on its surface, in contrast to the common negative charged surfaces found in most Trxs. This distinctive charge distribution is probably key to define initial encounter complexes that lead to a final specific interaction (21). In contrast, TrxT is more similar to other Trxs with respect to the charge distribution on its surface, but the fold itself is different. In this case, the presence of the C-terminal extension modulates the overall shape of this TrxT protein. This extension is connected to the protein core by a covalent loop, but even in the presence of this disulfide bond, the C-terminal region is flexible, as characterized by Nuclear Magnetic Resonance. When this extension is removed by introducing a premature stop codon, TrxR reduces the disulfide bond between Cys32 and Cys35 more efficiently than in the full-length protein but in parallel, the protein stability is decreased. Thus, this additional disulfide bond stabilizes a closed conformation that regulates the redox activity, hindering accessibility to the conserved catalytic site *in vitro*.

These two germline-specific Trx structures reveal new features of the proteins that can guide future work aimed at understanding embryo development and redox homeostasis in *Drosophila*. Moreover, due to their specific structural characteristics, they can help in the design of small molecular binders to modulate native redox homeostasis. These molecules can also provide new applications in the control of plagues that cause human diseases and/or that bring about economic losses by damaging crop production.

## MATERIAL AND METHODS

### Protein expression and purification

Dhd and TrxT sequences from *Drosophila melanogaster* were amplified from genomic DNA and cloned into the pOPINF expression vector. Constructs contained an N-terminal *His*10-tag followed by a 3C cleavage site. All clones were confirmed by DNA sequencing. All protein constructs were expressed in *E. coli* BL21 (DE3) Rosetta following standard procedures as previously described (22-24). Unlabeled samples were prepared using Luria Broth (LB) (Melford) and minimal media M9 with ^15^NH_4_Cl and/or D- [^13^C] glucose (Cambridge Isotope Laboratories, Inc) were used to prepare the labeled samples (25). Cells were cultured at 37 °C to reach an OD_600_ of 0.8-1.0. After induction with IPTG (final concentration of 0.5 mM) and overnight expression at 20 °C, bacterial cultures were centrifuged, and cells were lysed using an EmulsiFlex-C5 (Avestin) or a Vibra-Cell™ (Sonics) in the presence of Lysozyme and DNase I and in PBS buffer at pH 7.5. The soluble supernatants were purified by nickel-affinity chromatography (HiTrap Chelating HP column, GE Healthcare Life Science) using an NGC Quest 10 Plus Chromatography System (BIO-RAD). Eluted proteins were digested with 3C proteases at room temperature and further purified by size exclusion chromatography on HiLoadTM Superdex 75 16/60 prepgrade columns (GE Healthcare) equilibrated with 10 mM Tris-HCl pH 7.5, 100 mM NaCl. For crystallography, the last step of purification was performed using 20 mM Tris-HCl pH 7.5, 100 mM NaCl, and 2 mM ZnCl_2_. Both Trxs were stable and well folded in solution, as revealed by the NMR data. The 1D and 2D NMR data show a well-dispersed pattern of chemical shifts indicative of a folded sample (1H, 15N-Heteronuclear 2D Single Quantum Coherence, HSQC).

Purified proteins were verified by Liquid chromatography-Mass Spectrometry (LC-MS) using an ACQUITY UPLC Binary Sol MGR LC system (Waters) equipped with a BioSuite Phenyl 1000Å column (Waters, 10 μm RPC 2.0×75 mm) at a flow rate of 100 μL/min. The column outlet was directly connected to the mass spectrometer, which acquired full MS scans (400-4000 m/z) working in positive polarity mode. Samples were eluted using a linear gradient from 2% to 5% B in 5 min and from 5% to 80% B in 60 min (A= 0.1% Formic Acid, FA, in water, B= 0.1% FA in CH3CN) and analyzed using the MassLynx™ Software (V4.1.SCN704, Waters). The purity of the recombinant proteins was over 95%, as shown by the MS analysis.

### Sequence identification and clustering

Dhd and TrxT sequences are annotated in databases using different names. To avoid confusion, these sequences were retrieved using a protein BLAST (Basic Local Alignment Search Tool, *Blastp*) search in the NCBI Blast server, restricted to the non-redundant database and limited to dipteran insects. Using the guide trees generated with the alignments, we clustered the different Trx containing proteins and selected those clusters with TrxT and Dhd sequences for the analysis. Each cluster was manually inspected for outliers and re-aligned using Clustal Omega (26). Acession numbers and names are collected in Table 1 and Table 2. ESPript 3.0 (27) and BoxShade (https://embnet.vital-it.ch/software/BOX_form.html) were used to generate the figures as indicated in the figure legends. Secondary structure was predicted using the servers JPredv4 (28) and PrDOS (29).

**Table 1.** Data collection and refinement statistics.

|  | *Thioredoxin-T, TrxT* | *Deadhead, Dhd* |
| --- | --- | --- |
| **PDB entry** | **6z7o** | **6zmu** |
| Resolution range* | 35.13-2.24 (2.32-2.24) | 77.31-1.95 (2.05-1.97)<sup>#</sup> |
| Space group | P212121 | P43212 |
| Unit cell | 46.09 49.31 54.28 90 90 90 | 111.83 111.83 107 90 90 90 |
| Total reflections | 40618 (3817) | 473364 (21602) |
| Unique reflections | 4697 (457) | 43714 (2187) |
| Multiplicity | 8.0 (8.4) | 10.8 (9.9) |
| Completeness (%) | 99.94 (99.78) | 90 (38.3)<sup>#</sup> |
| Mean *I*/$\sigma$(*I*) | 9.37 (2.25) | 18.1 (1.5) |
| Wilson B-factor ($\text{\AA}^2$) | 39.06 | 44.68 |
| R-merge | 0.18 (0.86) | 0.07 (1.50) |
| R-meas | 0.19 (0.91) | 0.07 (1.57) |
| CC1/2 | 0.99 (0.85) | 1.00 (0.62) |
| R-work | 0.20 | 0.19 |
| R-free | 0.23 | 0.22 |
| Number of non-H atoms | 904 | 3606 |
| macromolecules | 872 | 3436 |
| ligands | 2 | 76 |
| Protein residues | 110 | 423 |
| R.m.s.d., bonds ($\text{\AA}^2$) | 0.010 | 0.86 |
| R.m.s.d., angles ($^\circ$) | 0.93 | 0.99 |
| Ramachandran favored (%) | 98.11 | 97.83 |
| Ramachandran allowed (%) | 1.89 | 2.17 |
| Ramachandran outliers (%) | 0 | 0 |
| Rotamer outliers (%) | 0 | 3.2 |
| Clash-score | 9.18 | 6.95 |
| Average B-factor ($\text{\AA}^2$) | 40.64 | 39.07 |
| macromolecules | 40.61 | 37.61 |
| ligands | 45.00 | 97.88 |
| solvent | 41.10 | 44.83 |

\*Values in parentheses are for the highest resolution shell. <sup>#</sup>Anisotropy correction by STARANISO/autoPROC with CC1/2 criterion used for the resolution cut-off (35),(36). Ellipsoidal completeness (%): 96.7 (70.6).

**Table 2.** Dhd acronyms and entries used in the protein alignment. BACDO, CERCA, MUSDO and ZEUCU entries were identified using Psi-Blast and are included as Dhd-family members based on the conservation of Arg/Lys residues characteristic of Dhd proteins. The identity and similarity to Dhd and Trx-2 DROME proteins are indicated to highlight that similarity cannot be exclusively used to classify Dhd proteins. Anopheles sequences (*Nematocera*) were also retrieved using Psi-Blast but in this case these proteins are considered to be Trx-2 proteins because the similarity/identity is observed for residues conserved between Trx-2 and Dhd but not for additional Arg/Lys residues only present in Dhd proteins. Calculated isoelectric points (59) are included to highlight that Dhd proteins have IP values larger than 7 and very often larger than 8. A sequence comparison of these divergent sequences is included as Supplementary Figure1F.

| Acronym | Species | RefSeq | Isoelectric Point |
| --- | --- | --- | --- |
| DROME | *Drosophila melanogaster* | NP_001284882.1 | 8.7 |
| DROYA | *Drosophila yakuba* | XP_002100027.1 | 8.7 |
| DROFI | *Drosophila ficusphila* | XP_017051920.1 | 8.6 |
| DROWI | *Drosophila willistoni* | XP_002071506.1 | 8.9 |
| DROSE | *Drosophila serrata* | XP_020803791.1 | 8.7 |
| DROVI | *Drosophila virilis* | XP_002057023.1 | 8.8 |
| DROGR | *Drosophila grimshawi* | XP_001992161.1 | 8.5 |
| DROBP | *Drosophila bipectinata* | XP_017090607.1 | 8.8 |
| DRONO | *Drosophila novamexicana* | XP_030568677.1 | 8.8 |
| DROTA | *Drosophila takahashii* | XP_017009012.1 | 8.7 |
| DROER | *Drosophila erecta* | XP_001976950.1 | 8.8 |
| DROAN | *Drosophila ananassae* | XP_014761224.1 | 8.8 |
| LUCCU | *Lucilia cuprina* | XP_023296875.1 | 8.8 |
| DROBI | *Drosophila biarmipes* | XP_016961519.1 | 8.7 |
| DRORH | *Drosophila rhopaloa* | XP_016983377.1 | 8.8 |
| DROSU | *Drosophila suzukii* | XP_016923010.1 | 8.9 |
| STOCA | *Stomoxys calcitrans* | XP_013114326.1 | 8.4 |
| DROKI | *Drosophila kikkawai* | XP_017030393.1 | 8.7 |
| DROEU | *Drosophila eugracilis* | XP_017083376.1 | 8.7 |
| SCALE | *Scaptodrosophila lebanonensis* | XP_030378843.1 | 8.9 |
| DROBU | *Drosophila busckii* | XP_017851482.1 | 8.5 |
| DROMI | *Drosophila miranda* | XP_017134982.1 | 8.7 |
| DROPS | *Drosophila pseudoobscura* | XP_001355409.1 | 8.7 |
| DROOB | *Drosophila obscura* | XP_022227514.1 | 8.7 |
| DRONA | *Drosophila navojoa* | XP_017965737.1 | 8.6 |
| DROHY | *Drosophila hydei* | XP_023162016.1 | 8.2 |
| DROPE | *Drosophila persimilis* | XP_002027085.1 | 8.7 |
| DRONV | *Drosophila novamexicana* | XP_030568678.1 | 7.4 |
| DROMO | *Drosophila mojavensis* | XP_002010046.1 | 8.5 |
| DROGU | *Drosophila guanche* | A0A3B0KTY9_DROGU | 8.7 |
| ZEUCU | *Zeugodacus cucurbitae* | A0A0A1WFG6_ZEUCU | 8.8 |
| CERCA | *Ceratitis capitata* | W8CBQ7_CERCA | 8.1 |
| MUSDO | *Musca domestica* | A0A1I8M261_MUSDO | 8.0 |
| BACDO | *Bactrocera dorsalis* | A0A034W4B0_BACDO | 8.8 |

| Acronym | Species | Dhd (IP 8.7) |  | Trx-2 (IP 4.6) |  | IP |
| --- | --- | --- | --- | --- | --- | --- |
|  |  | Identity | Similarity | Identity | Similarity |  |
| BACDO | *Bactrocera dorsalis* | 49.8% | 83.0% | 51.0% | 73.0% | 8.8 |
| CERCA | *Ceratitis capitata* | 54.0% | 72.0% | 48.0% | 75.0% | 8.1 |
| MUSDO | *Musca domestica* | 52.0% | 88.2% | 46.0% | 72.0% | 8.0 |
| ZEUCU | *Zeugodacus cucurbitae* | 43.0% | 71.0% | 60.0% | 75.0% | 8.8 |
| 9DIPT | *Anopheles braziliensis* | 40.8% | 82.5% | 61.0% | 80.0% | 4.8 |
| ANOFN | *Anopheles funestus* | 45.0% | 84.7% | 56.0% | 78.0% | 4.6 |

**Table 3.** TrxT protein acronyms and entries used in the alignments.

| Acronym | Species | RefSeq |
| --- | --- | --- |
| DROME | *Drosophila melanogaster* | Q8IFW4.1 |
| DROSE | *Drosophila sechellia* | XP_002036934.1 |
| DROSI | *Drosophila simulans* | XP_016038140.1 |
| DROMA | *Drosophila mauritiana* | XP_033171201.1 |
| DROBI | *Drosophila biarmipes* | XP_016962073.1 |
| DROSU | *Drosophila suzukii* | XP_016923019.1 |
| DROER | *Drosophila erecta* | XP_001976949.1 |
| DROYA | *Drosophila yakuba* | XP_002100028.1 |
| DROTA | *Drosophila takahashii* | XP_017009366.1 |
| DROAN | *Drosophila ananassae* | XP_001964015.1 |
| DROBP | *Drosophila bipectinata* | XP_017102470.1 |
| DROEU | *Drosophila eugracilis* | XP_017065989.1 |
| SCALE | *Scaptodrosophila lebanonensis* | XP_030378842.1 |
| DROWI | *Drosophila willistoni* | XP_002071507.1 |
| DROEL | *Drosophila elegans* | XP_017129970.1 |
| DROPE | *Drosophila persimilis* | XP_002021871.1 |
| DROMI | *Drosophila miranda* | XP_017134981.1 |
| DROPS | *Drosophila pseudoobscura* | XP_001355410.2 |
| DROOB | *Drosophila obscura* | XP_022227513.1 |
| DROVI | *Drosophila virilis* | XP_002057022.1 |
| DROBU | *Drosophila busckii* | XP_017850961.1 |
| DRONO | *Drosophila novamexicana* | XP_030568676.1 |
| DROHY | *Drosophila hydei* | XP_023162014.1 |
| DROGU | *Drosophila guanche* | SPP89275.1 |
| DROMO | *Drosophila mojavensis* | XP_002011512.1 |
| DRONA | *Drosophila navojoa* | XP_017963337.1 |
| DROAR | *Drosophila arizonae* | XP_017870321.1 |
| DROFI | *Drosophila ficusphila* | XP_017050179.1 |
| BACDO | *Bactrocera dorsalis* | XP_011199125.1 |
| ZEUCU | *Zeugodacus cucurbitae* | XP_011180011.1 |
| BACOL | *Bactrocera oleae* | XP_014095633.1 |
| RHAZE | *Rhagoletis zephyria* | XP_017484976.1 |
| CERCA | *Ceratitidis capitata* | XP_004521006.1 |

Isoelectric Point values are below 5

### NMR experiments

NMR data were recorded on a Bruker Avance III 600-MHz spectrometer equipped with a quadruple (^1^H, ^13^C, ^15^N, ^31^P) resonance cryogenic probe head and a z-pulse field gradient unit at 298 K.

Heteronuclear ^1^H–^15^N nuclear Overhauser enhancement (NOE), ^15^N -transverse relaxation time (T2) and 15N-longitudinal relaxation time (T1) spectra were acquired using pulse sequences described in the literature (30). ^1^H–^15^N NOE measurements were acquired using interleaved 2D-heteronuclear single quantum-correlation spectroscopy (HSQC), with and without ^1^H saturation, with 256 15N points and 24 (TrxT) or 40 (Deadhead) scans per t1 increment. ^1^H saturation was implemented using a 120° 1H pulse train with 5-ms intervals. For T2 measurements, a series of experiments was performed with nine relaxation delays (0, 17, 34, 51, 68, 102, 136, 170, and 238 ms) using the ^15^N Carr–Purcell–Meiboom– Gill pulse train (30).

For T1 measurements, a series of experiments was conducted with twelve relaxation delays (20, 50, 110, 160, 270, 430, 540, 700, 860, 1080, 1400 and 1720 ms). A series of ^1^H off-resonance 180° pulses was applied at 5-ms intervals to suppress cross correlation during the relaxation delay.

Raw data were processed and analyzed using the TopSpin 3.5 software (Bruker BioSpin). T1 and T2 relaxation times were calculated by non-linear least square fits of signal decays to an exponential decay function, S/S0=exp(-t/T1,2) (30). *τ*_c_ values were calculated using the Stokes equation (31):

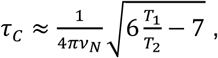

Where *v*_*N*_ is the ^15^N frequency in Hz.

^1^H–^15^N heteronuclear NOE values and ^15^N-relaxation rates (R1=T1-1 and R2=T2-1) were used to estimate the flexibility of the peptide.

Triple resonance experiments were acquired to obtain the backbone assignment of TrxT and Dhd using the *NMRlib 2.0* package (32). Specific proline backbone assignment was assisted with dedicated experiments (33). The data were processed using NMRPipe (34) and assigned using Cara (http://www.bionmr.com/).

### Crystallization

TrxT was concentrated to 15 mg/mL in 10 mM Tris pH 7.5, 100 mM NaCl, and 5 mM ZnCl_2_. Screenings and optimizations were prepared by mixing 100 nL of the protein solution and 100 nL of reservoir solution in 96-well plates. Crystals were grown by sitting-drop vapor diffusion at 20 °C. Crystals of TrxT were obtained in 15.0% w/v PEG 4000, 0.2 M potassium bromide. The Dhd sample was concentrated to 10 mg/mL in 20 mM Tris pH 7.2, 100 mM NaCl, and 5 mM TCEP. Crystals were grown by sitting-drop vapor diffusion at 20 °C. Screenings were prepared in 3 drop 96-well plates by mixing protein and reservoir solution at 2:1, 1:1 and 1:2 ratios, to final volumes of 300 nL. The best diffracting crystal was obtained in a drop containing 200 nL of protein sample and 100 nL of reservoir solution of 0.1 M sodium citrate pH 5.0, 3.2 M ammonium sulfate. It was cryoprotected by manual transfer to 0.1 M sodium citrate pH 5.0, 3.6 M ammonium sulfate.

### Data collection and structure determination

Diffraction data for TrxT were recorded on the beamlines ID23-1 and ID23-2 at the European Synchrotron Radiation Facility (ESRF) and for Dhd at the ALBA Synchrotron Light Facility (BL13-XALOC beamline), Barcelona, Spain. Diffraction data were processed with Mosflm and XDS, and scaled and merged with SCALA, either alone or with autoPROC (35). Anisotropy correction was applied using STARANISO (Dhd) (36). Initial phases were obtained by molecular replacement with Trx2 from *D. melanogaster* using PHASER (37,38) from the CCP4 suite (39) (search model PDB code: 1XWA). REFMAC and PHENIX (40,41) were used for the refinement, and COOT (42) for the manual improvement of the models. Figures were generated with COOT and UCSF Chimera (43).

### Differential Scanning Calorimetry

Experiments were performed in a StepOnePlus Real-time PCR System (Applied Biosystems). The assay was performed on 96-well plates (MicroAmp Fast 96-Well Reaction Plate, Applied Biosystems), in a total volume of 25 μL for each reaction. For stability screening, Slice pH™ HR2-070 (Hampton Research) was used. For additive screening (0-10 mM DTT, 0-10 mM TCEP, 0-10 mM ZnCl_2_, CaCl_2_, MgCl_2_), individual melting curves were acquired in triplicates and repeated twice. For each condition, the final protein concentration was 10 μM. SYPRO orange dye (sigma) was used at 60x dilution starting from a 5000x stock solution. Plates were sealed with optical quality sealing tape (Platemax). Samples were equilibrated for 60 s and analyzed using a linear gradient from 25°C to 95°C in increments of 1°C/min, recording the SYPRO orange fluorescence throughout the gradient.

### Thioredoxin Activity Assay

To test the activity of wild-type TrxT in comparison to shorter constructs without the C-terminal extension, a kit for assaying mammalian Trx1 using a 96-well microplate format was used (IMCO Corporation Ltd AB, FkTRX-02-V2). This method is based on the reduction of insulin disulfides by Trx with TrxR and NADPH as ultimate electron donors. The emission at 545 nm after excitation of eosin-labeled insulin at 520 nm was recorded for 30 min or up to 60 min. The assay was carried out using the following protocol. Briefly, Trx1 was diluted to a final concentration of 12 µg/ml (1 µM), and 5 µl of freshly prepared β-NADPH solution was added and incubated for 30 min. Before emission was recorded at 545 nm, 20 µl of the fluorescent substrate was added to all wells to start the reaction. The increasing fluorescence intensity for the time of the reaction within a linear range was calculated to obtain a standard curve for hTRX-1 activity. To test the activity of wild-type TrxT and short TrxT without the C-terminus (residue 1-111) each construct was freshly produced, and the concentration was set to 1 µM (determined by NanoDrop™ and controlled by SDS-Page). The assay was repeated three times, with two different batches of fresh protein.

## RESULTS

### Sequence comparison of *Drosophila* Trx proteins

A comparison of *Drosophila melanogaster* Trx sequences to vertebrate proteins revealed that the active site and the main features of the sequence are highly conserved. However, substantial differences are observed in the first strand and also in the second half of the protein (Supplementary Figure 1C). These regions are highly conserved in mammals but are more variable in other vertebrates and in *Drosophila melanogaster*. We expanded our analysis to include new Dhd and TrxT sequences deposited in databases after the analysis described in the literature in 2007 (20). Since the new proteins were not annotated as Dhd or TrxT, we used protein BLAST (Basic Local Alignment Search Tool, *Blastp*) to search in the NCBI database for new candidates. The search was first limited to highly conserved sequences using the *Drosophila melanogaster* TrxT sequence as the query. Using the guide trees generated with the alignments, we clustered and selected those groups that contain either Dm TrxT or Dhd sequences and re-aligned them using Clustal Omega (26). To identify distantly related protein sequences, we also used Psi-Blast and the EMBL-EBI search tools (44). Using these approaches, we found Dhd sequences belonging to *Schizophora* (true flies section), with representative sequences in the sub-sections *Calyptratae (Musca domestica)* and *Acalyptratae* (superfamilies *Ephydroidea and Tephritoidea)* with many entries of the genus *Drosophila.* The comparison of these sequences reveals the presence of conserved of abundant Lys and Arg residues (reflected by Isoelectric point values higher than 8), which are absent in the Trx-2 and TrxT sequences (Figure 1C, Supplementary Figure 1D and Table 2).

With respect to TrxT, the most obvious differences in these proteins are the presence of a third Cys residue at position 93 in sequences belonging to the subgenus *Sophophora* (specifically in the subgroups *Melanogaster* and *Suzukii* as well as in the subgroup *Pseudoobscura)*, and a highly divergent C-terminal domain. This domain is variable in length and sequence, ranging from 33 residues in *Bactrocera oleae* (olive fruit fly) up to 73 in *D. ananassae*. This domain often contains one or two additional Cys and many negatively charged residues, but lacks conserved hydrophobic residues (Figure 1D,E and Supplementary Figure 1E). In fact, in *Drosophila melanogaster* TrxT, secondary structure predictions of the C-terminal domain using JPredv4 (28) and the protein disorder prediction system (PrDos) (29) identified a short region with low disorder propensity surrounding the conserved Cys (Figure 1E,F).

As previously noticed (20), Trx-2 is the *Drosophila melanogaster* Trx with the highest sequence similarity to other potential hits detected in other insect species. This feature confirms that Trx-2 is probably the ancestral Trx protein in Diptera (20), and that TrxT and Dhd are the result of duplication events after the *Brachycera* and *Nematocera* separation.

### Biophysical characterization of Dhd and TrxT recombinant proteins

We expressed and purified both recombinant proteins and observed that Dhd and TrxT are stable at opposite pH values: TrxT is thermally stable with a T_m_ of > 70 °C in alkaline buffers, whereas acidic buffers are necessary for Dhd to reach the same stability (Figure 2A, Supplementary Table 2). Since the C-terminal domain was predicted to be unfolded, we expressed an additional construct without this region (TrxT-111). Already during purification, TrxT-111 formed disulfide-linked dimers, thereby confirming the propensity of the extra Cys93 to participate in disulfide bonds. Additionally, the thermal stability of this construct was reduced by approximately 20 °C with respect to the full-length protein. Its stability was unaffected by the addition of DTT, which should reduce the dimer. Upon addition of DTT to full-length TrxT, the T_m_ value was reduced by approximately 20° C (Figure 2B), indicating that the presence of the C-terminal domain increases the stability of the protein, perhaps at the expense of a second intra-molecular disulfide. The addition of DTT to Dhd did not affect the thermal stability of the protein (Supplementary Table 2).

**Figure 2.**
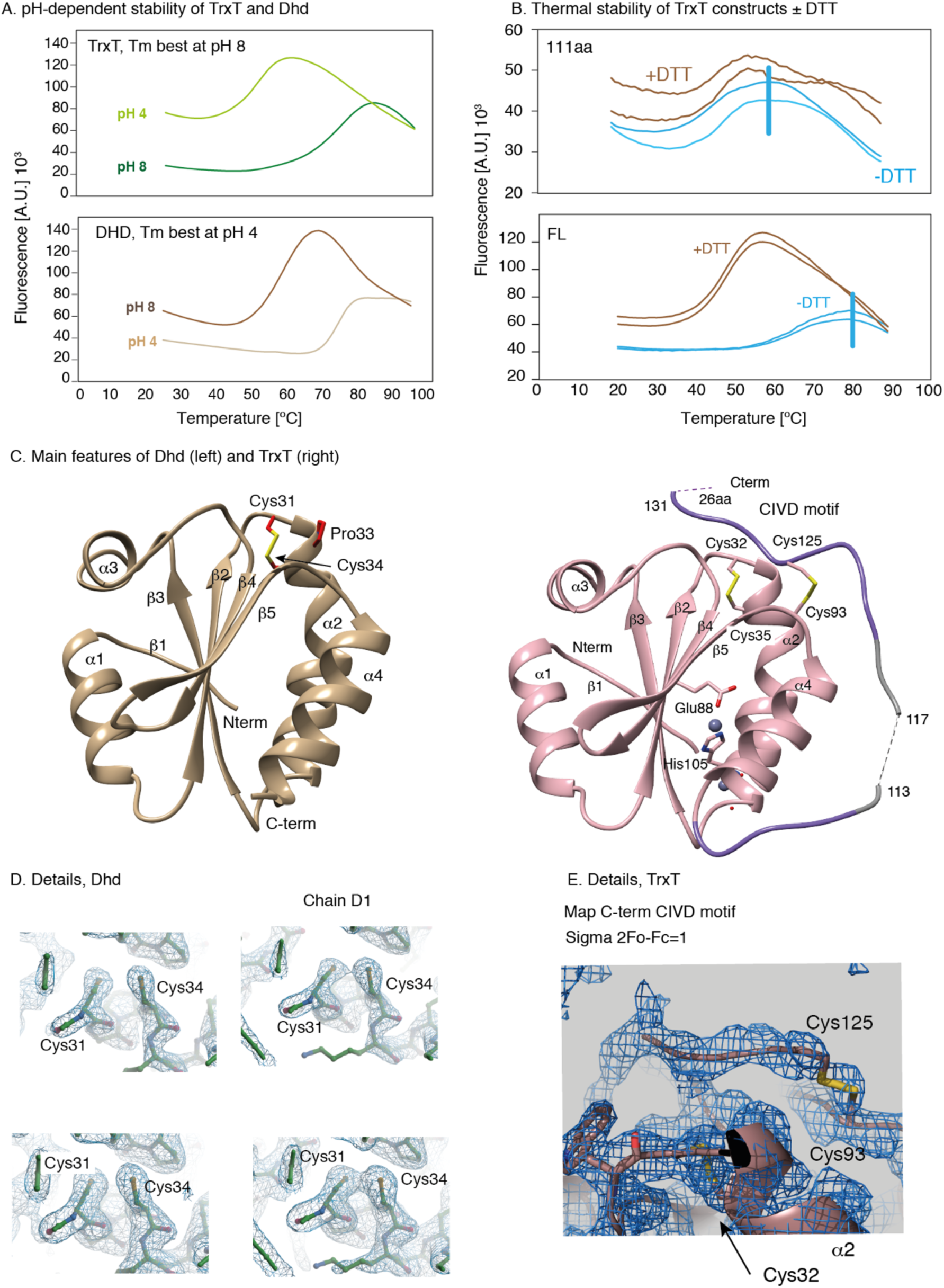
Crystal structures of *D. Melanogaster* Dhd and TrxT proteins. **A.** Thermal shift assay of TrxT and Dhd after incubation of natively folded proteins with SYPRO Orange dye in a 96-well PCR plate, at two different pH values. As the proteins unfold with the temperature, the SYPRO Orange fluorescence emission increases. TrxT is thermally stable with a T_m_ of > 70° C in alkaline buffers, whereas acidic buffers are necessary for Dhd to reach the same stability. Values were obtained as triplicates and were collected at different conditions (Supplementary Table 2). **B.** Thermal stability of TrxT FL and TrxT-111aa constructs in the presence or absence of DTT (duplicates). TrxT FL is ∼20 ° C more stable than the protein core. **C.** Cartoon representation of the crystal structures of Dhd protein (left) and TrxT (right) showing shows the oxidized forms. Cysteines 32 and 35 and Proline 34 in the catalytic motif are labeled. Secondary structure elements are also indicated. For TrxT, part of the C-terminal domain was connected to the Trx core domain by a disulfide bridge between Cys 93 and 125. **D.** Electron density map of active center forming Cys32 and Cys35. Left figures: reduced chain B. Right figures: partially oxidized chain D. Top 2Fo-Fc electron density map contoured at 2.0 sigma, bottom at 1.0 sigma. **E.** The Fo-Fc OMIT electron-density map for the bound C-terminal fragment contoured at 3.0 sigma (top) and the 2Fo-Fc map contoured at 1.0 sigma (bottom).

We also determined the behavior of the proteins in solution using NMR. To this end, TrxT and Dhd were additionally expressed as ^15^N and ^13^C and ^15^N labeled samples. Both proteins were well-folded using a 2D ^1^H-^15^N HSQC. However, while Dhd had well-dispersed amide resonances, TrxT displayed a mix of dispersed and overlapped resonances in the HSQC experiment, revealing a substantial conformational plasticity for about 50 residues, which probably correspond to the C-terminal domain (Supplementary Figure 2A). The overall T1 and T2 values (T1: 800 ms, T2:67 ms) and correlation times, *τ*_c_ =10 ns, of TrxT were independent of the presence or absence of DTT. These values agree with a monomeric ∼17 kDa protein (theoretical MW =17.5 kDa). For Dhd, we observed a similar monomeric behavior unaffected by the presence of DTT. In this case, we obtained a *τ*_c_ of 6.7 ns (T1:620 ms and T2:110 ms), which fits well with a monomeric protein of ∼11 kDa (theoretical MW =11.2 kDa).

### Structures of Dhd and TrxT

We used X-ray crystallography for the structural characterization of full-length Dhd and TrxT (Table 1). Attempts to obtain crystals were performed in the presence or absence of reducing agents (TCEP). However, TrxT crystals were only obtained in the absence of TCEP whereas the best Dhd crystals were obtained in buffers containing TCEP.

The asymmetric units (ASU) of TrxT and Dhd contain one and four monomers, respectively (Supplementary Figure 2B). In both structures, the core (residues 1-106 and 2-108 respectively) showed an alpha/beta fold that contains four central beta strands surrounded by four alpha helices. In these structures, the β1-strand has a single set of hydrogen bonds with the third strand oriented as two parallel strands, and this feature explains why it is not detected as a proper strand by visualization/analysis programs like Chimera. The β-strands of the C-terminal region (β2, β4 and β5) run anti-parallel but β2 and β3 are again oriented parallel to one another (Figure 2C). In agreement with previously determined structures, the α2 helices are slightly curved due to the bend caused by the presence of a Pro (TrxT) or a Ser (Dhd) residue in the middle part of the helix. The active sites (Cys-Gly-Pro-Cys) are located between the β2 strand and the N-terminal part of the α2 helix, as observed in all Trxs. The active sites are oxidized in the case of TrxT and reduced in the case of Dhd, although the electron density for one of the four monomers (chain D), indicates the presence of the catalytic Cys-Cys disulfide bridge in equilibrium with the reduced form (Figure 2C, D). In the case of TrxT, the crystals contain two TrxT molecules engaged with a symmetry-related neighbor stabilized through the coordination of a Zn atom (present in the crystallization condition). The Zn is bound between Asp65 and Glu69 from monomer A, and His105 and Glu88 from the monomer B (Supplementary Figure 2C).

When superimposed to the previously determined Trx-2 structure, the canonical fold of Dhd and TrxT proteins is highly conserved with minor differences observed for residues surrounding the catalytic site, especially when reduced and oxidized forms are compared (Figure 2D). The RMSD values calculated with respect to Trx-2 are ∼0.9 Å for Dhd and ∼1.2 Å for TrxT (for all residues in the core, including loops). When compared to other structures, one of the differences is found at P76, which in Dhd, TrxT and Trx-2 adopts a *trans* configuration, and not *cis*, as reported for other Trxs (2).

Moreover, we found that Cys93 in TrxT, located at the loop connecting β5 and α4, forms a covalent bond with Cys125 of the C-terminal domain, explaining the monomeric behavior of the protein in solution and in crystals. The density 2F_o_-F_c_ plotted at sigma 1 showed that the bound C-terminal fragment (Cys125-Asp128) is not present in all molecules in the crystal, with an occupancy of approximately 70% (Figure 2E). In addition, the regions connecting Gly107 to Asp111 and His120 to Cys125 were also traceable, but unfortunately, the small fragment connecting Asp111 to His120 and the last 26 aa were not visible in the density. These results indicate that the C-terminal domain contributes to the structure of the protein through the presence of a closed conformation, which prevents oligomerization via disulfide bonds between monomers. Since the crystal structures could not describe the full-length C-terminal domain, we considered the possibility of a certain degree of flexibility in this part of the protein, in agreement with the secondary structure predictions for this domain and with the NMR data characterized in solution (Supplementary Figure 2A).

### Dhd surface is positively charged

Eukaryotic Trxs frequently have negatively charged patches on their surfaces, as is the case of Dm Trx-2 (Supplementary Figure 1A) and TrxT (Figure 3A, left). However, *Drosophila melanogaster* Dhd does not follow this rule and presents an unusual positively charged surface (Figure 3A, right). The presence of positively charged patches is often taken as an indication of membrane binding to phospholipids or as a protein-DNA/RNA binding patch. In fact, known Dhd targets include protamine proteins, ribosomes and ribosome-associated factors, thereby suggesting a role of this specific charge distribution in selecting protein partners (18). The residues responsible for these patches are highly conserved in other *Schizophora* sequences (Figure 1C, Supplementary Figure 1D). To illustrate this conservation, we have modeled six Dhd sequences onto the *Drosophila melanogaster* Dhd structure. These models reflect that the positively charged patches are likely to be present in other Dhd proteins (Supplementary Figure 4A,B), indicating that positively charged patches are a common feature distinguishing Dhd from other Trxs.

**Figure 3.**
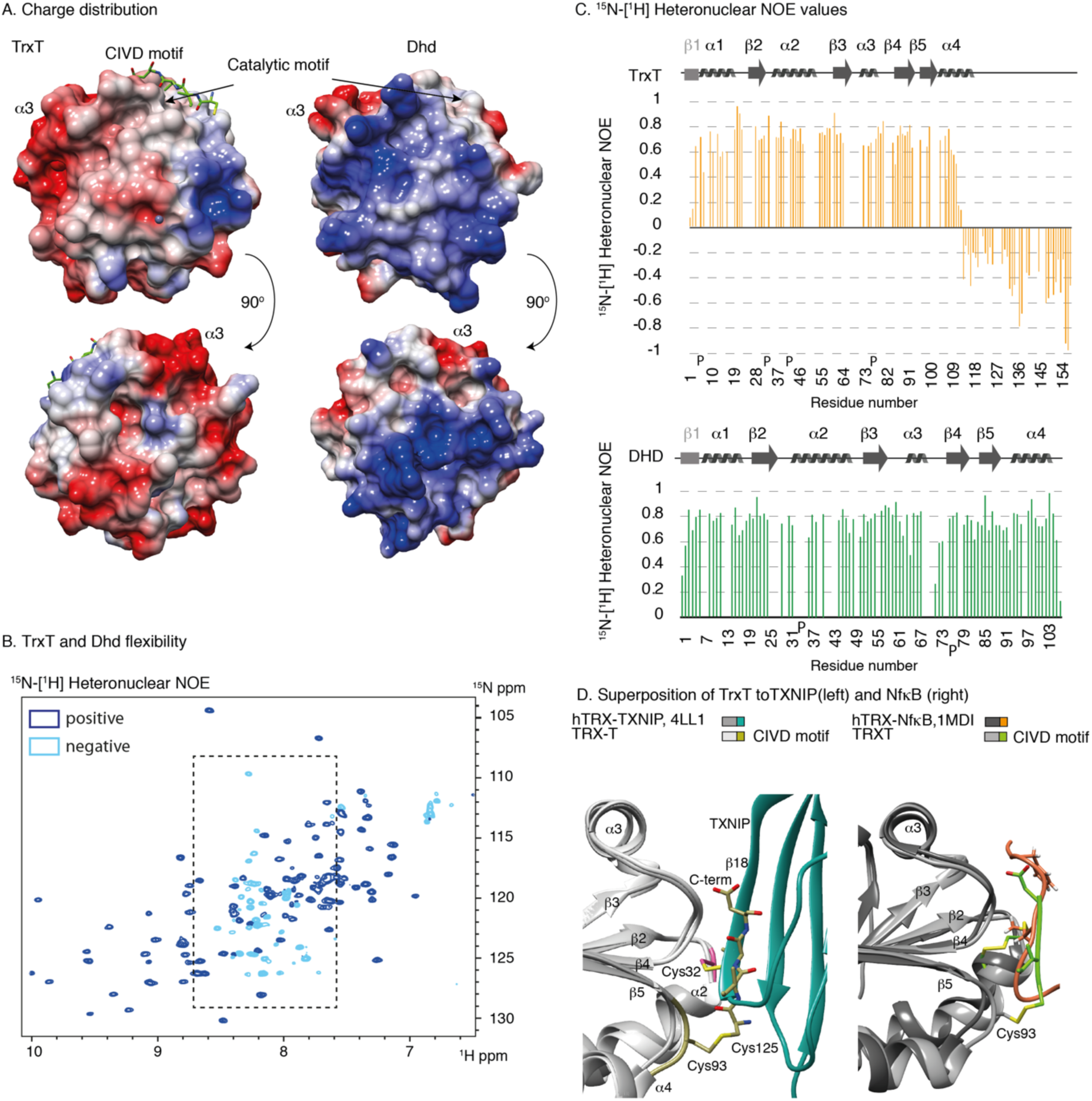
Charge distribution, flexibility and comparison to other Trx protein complexes. **A.** Charge distribution for the TrxT protein (left) and Dhd (right). TrxT shows a typical surface charge for a thioredoxin domain whereas Dhd shows unusual positively charged patches. **B.** Comparison of the overall structure of the TrxT and TXNIP (left, PDB:4LL1) and NFκB (right, PDB:1MDI) using human Trx for the fitting. The C-term CIVD motif of TrxT (shown in chartreuse) occupies the same place as the Trx partners in the human Trx complex structures. **C.** ^15^N-[^1^H] Heteronuclear NOEs of TrxT in the absence of DTT. Assignments corresponding to the flexible residues (all located at the C-terminal domain) are shown in Supplementary Figure 3A. The same experiment with DTT is shown as Supplementary Figure 3B. Equivalent experiments corresponding to Dhd are shown as Supplementary Figure 3C. **D.** ^15^N-[^1^H] Heteronuclear NOEs were measured as duplicates for TrxT (orange) and Dhd (green). Negative values are characteristic of highly flexible regions. The missing bars correspond to prolines and the residues for which amide resonances were not assigned.

### NMR relaxation experiments confirm that the TrxT C-terminal extension is flexible

Flexible regions have favorable relaxation properties and give rise to good signal-to-noise NMR backbone resonance data, provided that signals are not highly overlapped. In this case, to obtain the sequence-specific resonance assignment, we had to combine backbone triple resonance experiments (32),(33) with site-specific amino acid-type information using iHADAMAC experiments due to amino acid repetitions in the sequence (45). A 3D ^15^N NOESY-HSQC spectrum was used to validate assignments for the secondary structural elements of TrxT. The absence of sequential and medium-range NOEs for residues located at the C-terminal domain indicate that this region is highly flexible when compared to the rest of the protein. With this approach, we identified 123 residues of the 157 present in TrxT, including 75/101 residues in the protein core, as well as 48/51 residues in the flexible C-terminal domain.

To identify the residues showing rapid motions, we determined ^15^N-[^1^H] heteronuclear NOE values in solution, since residues in random coil regions typically exhibit faster internal motions than in structured regions. Changes in the redox state of Cys also affect the ^13^Cβ chemical shift values (CSV), being between 26 and 38 ppm for reduced/oxidized cysteines in α-helices, and between 30 and 43 ppm for β-strands (46). In TrxT, Cys32 and Cys35 have Cβ CSV corresponding to oxidized forms (located at helix 2), whereas Cys93 and Cys125 (both located at loops) are 30.4 and 34.2 ppm, respectively. These values indicate a mixture of redox states and conformations, as previously observed in the crystals, where the Cys93-Cys125 disulfide is present in ∼75% of the conformations. Moreover, we observed intense backbone signals for residues 108-157, characteristic of residues affected by rapid motions in the presence or absence of DTT. In both conditions, negative NOE values were detected for the most C-terminal part of the domain (135-157 residues), suggesting fast motions for this region and the presence of an ensemble of conformations in rapid interchange. This region contains the highest ratio of charged residues (55%). Residues from Ala 106 up to His120 also displayed significantly lower NOE values, indicating intermediate to fast internal motions, faster than other residues assigned to well-defined secondary structure, thus corroborating the flexibility detected in crystals. The addition of DTT affects the resonances of residues located in this region, suggesting that the reduction of the Cys93-Cy125 bond enhances the overall flexibility of this domain (Figure 3B,C and Supplementary Figure 3A,B). These properties indicate that the C-terminal domain is able to experience rapid motions, sampling different conformations in solution even in the presence of a disulfide bond.

Backbone triple resonance experiments and ^15^N-[^1^H] heteronuclear NOE experiments were also performed for Dhd for comparison purposes. In this case, we identified 93 residues of the 107 present in Dhd and Cys31 and Cys34 have Cβ CSV corresponding to reduced forms. All the ^15^N-[^1^H] heteronuclear NOE peaks were positive, thereby indicating that the Dhd fold is highly compact, as observed in the crystals (Figure 3C and Supplementary Figure 3C).

### The C-terminal domain of TrxT modulates the access to the catalytic site

Due to the orientation of the C-terminal fragment near the catalytic site, we hypothesize that this closed conformation affects the redox activity of the *Drosophila melanogaster* TrxT protein. To illustrate this hypothesis, we compared the structure of TrxT to that of other Trxs in complex with targets described in the literature. Overlaying the TrxT structure with that of human Trx in complex with the Thioredoxin-Interacting Protein (TXNIP, PDB: 4LL1) showed that the C-terminal fragment of TrxT binds in a similar manner and within the same site, as observed in the TXNIP-Trx complex (47). In fact, the TXNIP-Trx interaction serves to inhibit Trx redox activity by impairing access to the Trx catalytic site (48). Although there is a clear parallelism, the interaction involves distinct Cys residues: an intramolecular disulfide Cys93-Cys125 in TrxT versus the intermolecular disulfide between hTrx and TXNIP (Figure 3D, left). A similar interaction is also observed in the human Trx complex with one of its substrates, the transcription factor NfκB, bound to the catalytic Cys32 (PDB: 1MDI) (Figure 3D, right) (49). These structural similarities suggest a structural/functional convergence to provide a regulatory redox mechanism in TrxTs.

### The C-terminal domain of TrxT modulates its redox activity *in vitro*

In all organisms, the major function of TrxR is the maintenance of Trx in its reduced state (50). Most TrxRs are homodimeric enzymes whose catalytic site is located at the flexible C-terminus (1). TrxR activity is conferred by either a Cys residue in insects or by a Seleno-cysteine in mammals (51-53). Assuming that TrxT interacts with TrxR in a similar manner to that observed in the complex of human TrxR-Trx1, the presence of the C-terminal domain might affect the access to the catalytic site (54). Using the human Trx-TrxR complex as a template (PDB code: 3QFA), we docked Dhd or Trx-2 proteins onto the TrxR surface places. In both models, Cys32 site is accessible to the reductase, as observed in the human complex (Figure 4A, B). However, docking TrxT in a similar manner (with the bound C-terminal domain, as observed in the crystals) blocks the approximation of TrxR to Cys32 (Figure 4C). Since the catalytic loop of TrxR is flexible (54), we propose that a slight reorientation of this loop would permit a subtle variation of the mechanism, first reducing the Cys93-Cys125 bond—and promoting a transition from the closed to an open conformation—and then reducing the now accessible Cys32-Cys35 (Figure 4C,D). Of note, in humans, the reduction of non-catalytic disulfide bonds present in Trxs is normally carried out by GR proteins, but these proteins are absent in *Drosophila* species.

**Figure 4.**
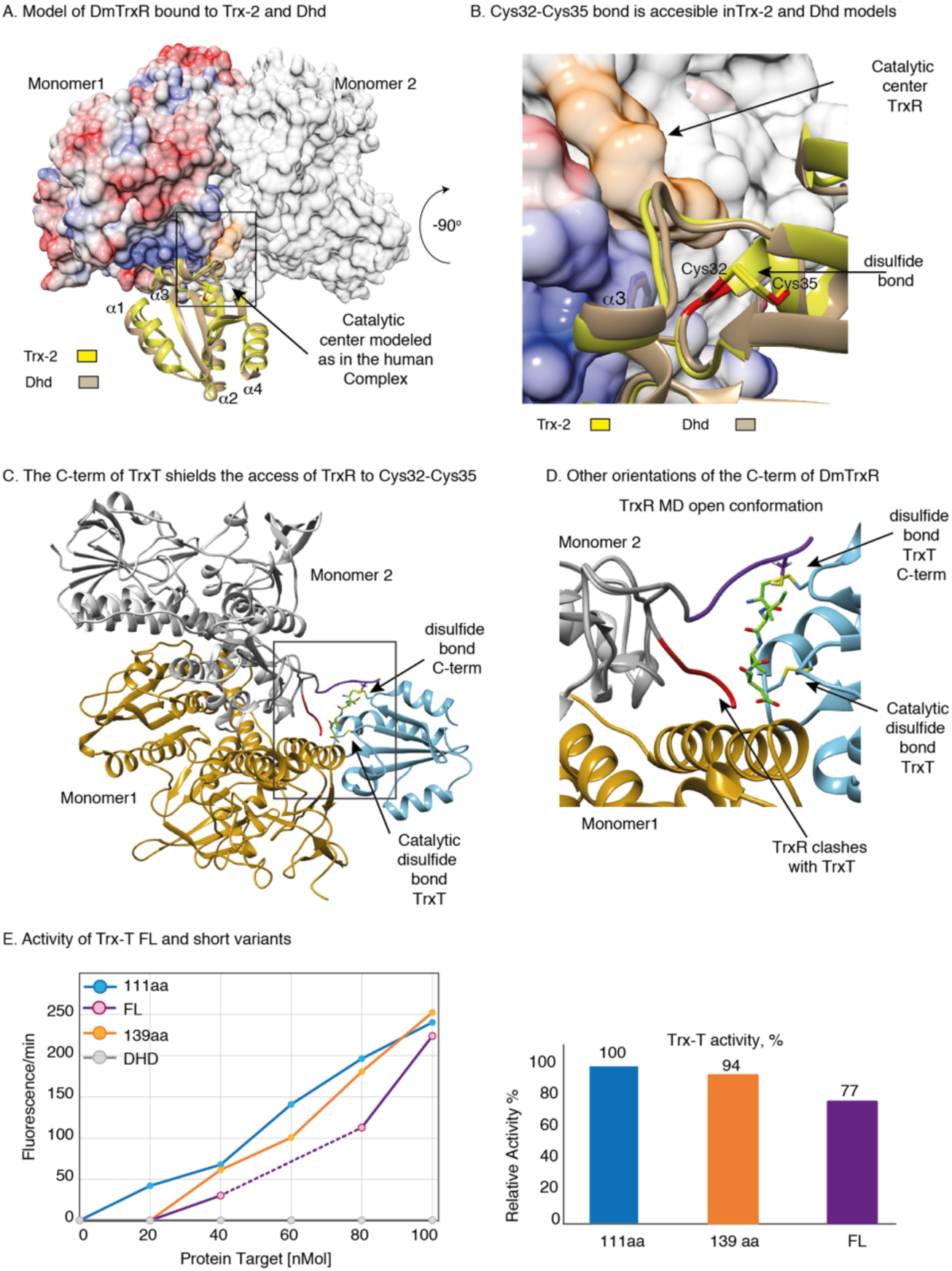
The C-term fragment modulates the stability and redox activity of TrxT. **A.** Model of Trx-2 and Dhd structures docked to Dm-TrxR as observed in the human TrxR-Trx complex. The catalytic center of TrxR is able to access the C32-C35 bond in both *Drosophila* thioredoxins. **B.** Close-up view of the interaction between the catalytic center of the reductase (colored in orange and indicated with an arrow) and the oxidized forms of Trx-2 (in yellow) and Dhd (in tan) 90° rotated with respect to the view shown in **C**. The catalytic Trx and Dhd Cys are shown in red. The α3 helix of Dm-Trx and Dhd that participates in direct contacts with the reductase is labeled. **C.** Model of TrxT docked to Dm-TrxR as depicted in Figure 4A. The catalytic center of TrxR as determined in the human complex (PDB:3QFA, shown in red) cannot access the C32-C35 bond in TrxT due to the presence of the C-term motif attached to Cys93 in the core domain (both sites are indicated with arrows). The orientation shown in purple will allow the reduction of Cys93-Cys125 of the tail. Once this step is achieved, a second reaction can occur to reduce the C32-C35 bond. **D.** Close-up view of the potential interaction between the catalytic center of the reductase (colored in red and purple and indicated with arrows) and the oxidized forms of TrxT (in blue), 90° rotated with respect to the view shown in **C**. The catalytic Trx and Dhd Cys are shown in red. The α3 helix of Dm-Trx and Dhd that participates in direct contacts with the reductase is labeled. **E.** Redox activity of TrxT constructs against an eosin-labeled insulin-S-S substrate using a commercial kit developed by IMCO corporation Ltd. The activity of the FL TrxT protein is slightly less than that of the totally processed C-term domain. However, the C-terminal domain contributes to the stability of the FL protein, increasing the melting temperature by ∼20 °C.

To evaluate our hypothesis, we analyzed the activity of two TrxT constructs (full-length, and 111aa) towards an eosin-labeled insulin peptide using a commercial assay, which provides a mammalian TrxR and a human Trx as a positive control. Even using the human TrxR, which has a smaller reduction potential than that of *Drosophila* (54), we observed that the full-length TrxT construct reduced the eosin-labeled insulin substrate by approximately 80% compared to the short construct lacking the C-terminal extension. No effect was observed for Dhd under these experimental conditions, probably due to the basic pH recommended for the assay, which compromises the stability of Dhd (Figure 4E). Overall, these results underline the hypothesis that the C-terminal domain shields TrxT active site, and modulates its redox activity.

## DISCUSSION

### Common and specific features of Dhd and TrxT structures

The Trx fold is defined by a core structure that contains three α-helices and four β-strands although most Trx structures fold as four α-helices and five β-strands (2). *Drosophila melanogaster* Dhd, TrxT and Trx-2 (4) lie in the middle of these two characteristic folds, displaying four α-helices and four β-strands (β2-β5), with the β1-strand being poorly defined. Apart from these differences at the secondary structure level, the overall 3D structure is well conserved with respect to other Trxs, including the presence of the catalytic site located at the N-terminal of helix α2. Being at its N-terminal part, the site is affected by the general positive dipole moment of α helices. This feature stabilizes the thiolate form and enhances its catalytic capacity, which decreases the pKa of Cys32. Another conserved characteristic is the presence of a negatively charged surface patch in helix α3, which is required to interact with the TrxR to recover the redox equilibrium.

Apart from these similarities, two main differences are observed in the structures of *Drosophila melanogaster* Dhd and TrxT with respect to other Trx structures. Whereas the Trx surfaces including TrxT are negatively charged, Dhd has extended positively charged patches. These areas are likely to be present in other structures of *Schizophora* species, as the Arg and Lys residues responsible for these patches are abundant and conserved in all Dhd proteins (Supplementary Figure 1D, F). When Dhd and Trx-2 sequences are compared, we noticed that the additional positively charged residues cluster at sites where Trx-2 sequences have negatively charged residues or hydrophobic amino acids. By displaying a positively charge distribution (and not negative as most Trxs), Dhd proteins would speed-up the selection of specific targets during initial encounter complexes of redox proteins *in vivo* (21,55). Among these redox reactions are the reduction of intermolecular disulfide bonds in *Drosophila* protamine oligomers to facilitate their eviction from DNA (14). To perform this function, Dhd proteins must get close to the protamine oligomers bound to DNA, and this approximation would be facilitated by charge complementarity (negative at the DNA backbone and positive at the Dhd surface). The charge complementarity would also favor the interaction with ribosomes and associated factors like NO66, which is a lysine-specific demethylase that removes methyl groups from histone H3, and with the ribosome protein 8 (Rpl8) hydroxylase (56).

The second difference observed between previously characterized Trxs and TrxT is related to the protein length. TrxT is composed of a highly-conserved core and a variable C-terminal domain, the latter being absent in most canonical Trxs. This C-terminal domain is mostly unstructured, with a high ratio of charged residues, and it is attached to the protein core via a disulfide bond in *Drosophila melanogaster*. This covalent link between Cys93 and Cys125 stabilizes a closed conformation that partially covers the catalytic site. We observed that many TrxT proteins have a Cys at the C-terminal domain but only a third of them have Cys93. The latter feature indicates that not all dipteran TrxTs can form a second disulfide bond as observed in *Drosophila melanogaster* and perhaps the C-terminal domains of other TrxT sequences might adopt different orientations. The high sequence variability of this region (almost species-specific) might also contribute as an additional switch to regulate TrxT-protein interactions.

Apart from the catalytic site, the presence of additional disulfide bonds has been observed in other Trxs, as for example in human Trx1. In this case, this bond involves Cys62 and Cys69 and inactivates the redox capacity of the protein (57). Compared to this inhibitory role in human Trx1, the formation of the additional disulfide bond in TrxTs seems to have a mild effect on their function and might provide a mechanism through which to increase the stability of the protein not only *in vitro*—as we have characterized—but perhaps also *in vivo*.

### Potential applications to guide structure-based drug design

Many drug-discovery strategies for aging, anti-cancer and Parkinson therapies (58) have taken advantage of model organisms like *Drosophila* as cost-effective alternatives to mammalian cellular/animal systems. This approach is based on the conservation of many pathways in metazoans as well as on a good knowledge of the differences. Having the structural knowledge at hand would facilitate the process of target identification and virtual/experimental screening of potential ligands and may also guide the docking of protein partners described in the literature (18). This knowledge will help to correlate binding effects with phenotypes during oxidative stress and redox signaling. In this context, the charge distribution of Dhd should drive the selection of molecular binders, these being different from those preferentially selected by Trx and TrxT counterparts. Moreover, the specific features displayed by TrxT and Dhd should be considered when screening molecules as binders to regulate human redox systems.

In addition to using *Drosophila* as a model for human diseases, our results may have applications for the design of inhibitory molecules to reduce and control fly plagues by selecting the germline Trx proteins as targets. These plagues, as black fly species that spread diseases such as river blindness in Africa and the Americas (World Health Organization), have an impact on human health. Others negatively affect the economy of many countries due to the losses in fruit and vegetable production worldwide.

## DATA AVAILABILITY

Atomic coordinates and electronic densities for the reported crystal structures have been deposited with the Protein Data bank under accession numbers 6ZMU (Dhd) and 6Z7O (TrxT).

## SUPPLEMENTARY DATA

Supplementary Data are available at NAR online.

## Author Contributions

R.F., E.A., B.B., R.P. and M.J.M. designed and performed most of the experiments and coordinated collaborations with other authors. E.A. cloned, expressed and purified all the proteins. E.A., P.M.M., M.C. and M.J.M. performed the NMR measurements and analyzed the data. R.F., E.A., B.B., and R.P. screened the crystallization conditions, collected X-ray data and determined the structures. R.F. B.B. R.P. and M.J.M. analyzed the structures. R.F and L.R. performed the thermal stability studies and tested the redox activity. All authors contributed ideas to the project. R.F., E.A. C.G. and M.J.M. conceived and supervised the project. M.J.M. wrote the manuscript with contributions from all other authors.

## ACKNOWLEDGEMENTS

We thank Dr. N. Berrow (IRB Barcelona, Protein Expression Unit) for help with some DNA constructs, protein purification and reagents, S. Llamazares for providing us with *Drosophila* DNA, C.M for support with some biochemical assays, and Dr. E. Guca for help with protein crystallization and data acquisition. We also thank the staff at the Automated Crystallography Platform (IRB Barcelona-CSIC) and at the ESRF (Grenoble) and ALBA synchrotrons (Barcelona) for access to the beamlines.

## FUNDING

R.F., R.P., and B.B. are co-funded by the European Union’s Horizon 2020 Research and Innovation Programme under the Marie Skłodowska-Curie COFUND actions of the IRB Barcelona, PROBIST and PREBIST Postdoc and Predoc Programmes (agreements IRBPostPro2.0_600404 and PROBIST_754510, and PREBIST_754558). C.G. and M.J.M are ICREA Programme Investigators. This work was supported by IRB Barcelona and by ERC AdG 2011 294603 and by PGC2018-097372-B-100 funded by ERDF/Ministry of Science, Innovation and Universities-Spanish State Research Agency (T.G.). Access to ALBA and to the ERSF synchrotrons was granted through BAG proposals 2018092972 and MX-1941, respectively. We gratefully acknowledge institutional funding from the CERCA Programme of the Catalan Government and from the Spanish Ministry of Economy, Industry and Competitiveness (MINECO) through the Centers of Excellence Severo Ochoa Award. Funding for open access charge: IRB Barcelona and the Marie Skłodowska-Curie COFUND actions.

## CONFLICT OF INTEREST

The authors declare no conflict of interest.

## TABLE AND FIGURES LEGENDS

**Supplementary Table 1.** Thioredoxin acronyms used in S-Figure 1.

| Alignment name | Species | RefSeq |
| --- | --- | --- |
| Trx1 - Human | *Homo sapiens* | AF313911.1 |
| Trx1- Marmoset | *Callithrix jacchus* | AF353204.1 |
| Trx1 - Mouse | *Mus musculus* | NP_035790 |
| Trx1 - Blind mole rat | *Nannospalax galili* | XP_008852851 |
| Trx1 - Horse | *Equus caballus* | NP_001075282 |
| Trx1 - Arabian camel | *Camelus dromedarius* | XP_010977020 |
| Trx - Sperm whale | *Physeter catodon* | XP_007130171 |
| Trx - Dingo | *Canis lupus dingo* | XP_025288383 |
| Trx - European Hedgehog | *Erinaceus europaeus* | XP_016049782 |
| Trx - Egyptian fruit bat | *Rousettus aegyptiacus* | XP_015985530 |
| Trx - Amur tiger | *Panthera tigris altaica* | XP_015393030 |
| Trx - White Rhinoceros | *Ceratotherium simum simum* | XP_004423376 |
| Trx- Chicken | *Gallus gallus* | NP_990784 |
| Trx - Giant devil catfish | *Bagarius yarrelli* | TSK22521 |
| Trx - Northern pike Lucius | *Esox lucius* | NP_001290895 |
| Trx - Torafugu | *Takifugu rubripes* | XP_003978603 |
| Trx - Xenopus Tropicalis A | *Xenopus tropicalis* | XP_002940306 |
| Trx - Xenopus Tropicalis B | *Xenopus tropicalis* | XP_031746215 |
| TrxT - D. melanogaster | *Drosophila melanogaster* | AAF46018 |
| Trx2 - D. melanogaster | *Drosophila melanogaster* | AAN10700 |
| Trx1/Dhd - D. melanogaster | *Drosophila melanogaster* | AHN59353 |

**Supplementary Table 2.** Tm values at different pH and buffers.

|  | Citrate | Acetate | Cacodylate | BIS-TRIS | ADA | MOPS | Tris | BICINE | AMPD |
| --- | --- | --- | --- | --- | --- | --- | --- | --- | --- |
| pH | 4.5 | 4.9 | 6.1 | 6.6 | 7.0 | 7.4 | 8.5 | 8.7 | 9.3 |
| TrxT | 52 ±0.2 | 62±0.6 | 54±0.3 | 65±0.2 | 68±0.1 | 69±0.1 | 68±0.1 | 65±0.1 | 54±0.2 |
| Dhd | 85±0.1 | 75±0.1 | 73±0.2 | 68±0.3 | 73±0.1 | 63±0.1 | 60±0.1 | 61±0.1 | 62±0.6 |

Citrate: Sodium Citrate tribasic di-hydrate, acetate: sodium acetate trihydrate, cacodylate: sodium cacodylate trihydrate, ADA, N-(2-acetamido) iminodiacetic acid, MOPS (3-(N-morpholino) propanesulfonic acid), Tris: trisaminomethane, BICINE: N,N-Bis(2-hydroxyethyl)glycine, AMPD (2-amino-2-methyl-1,3-propanediol).

**Supplementary Figure 1.**
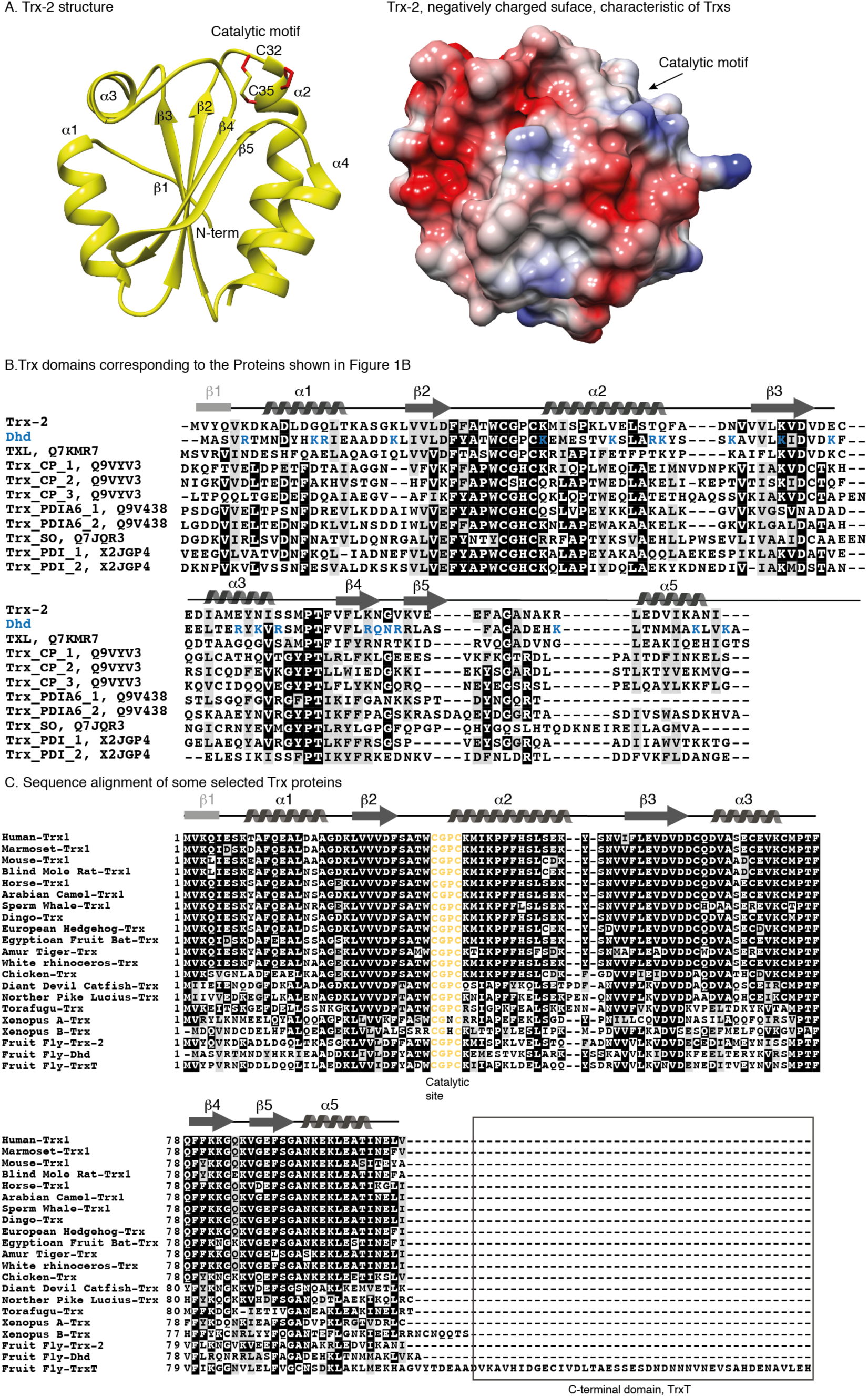
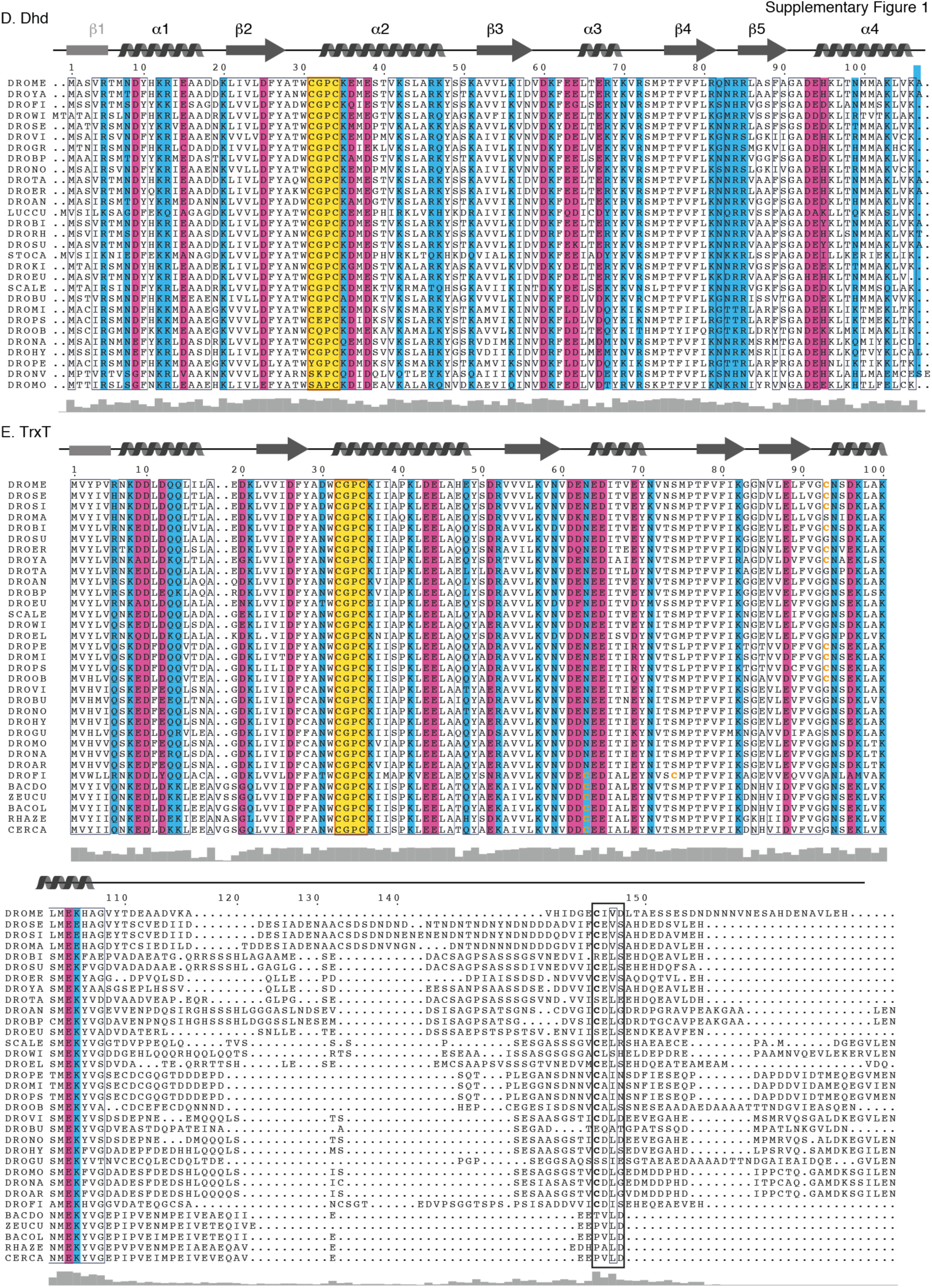
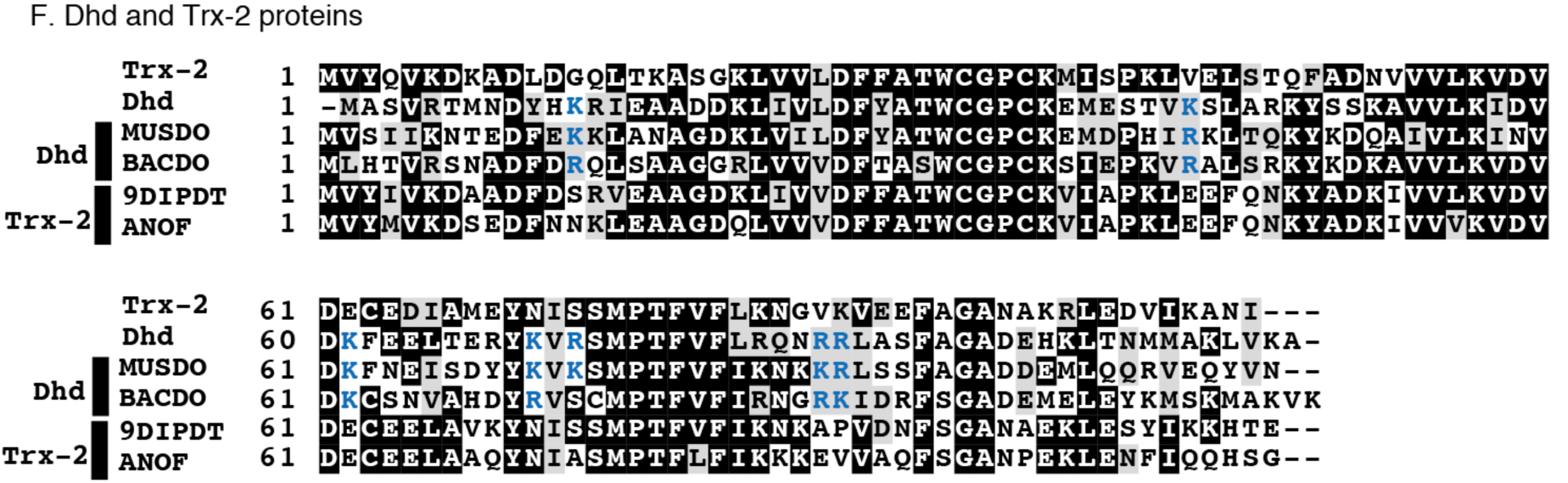
**A.** Ribbon diagram of the Trx-2 structure in its oxidized form (PDB:1XWA). All secondary structure elements are labeled. Molecular surface properties. Electrostatic potential of Trx-2, in which positively and negatively charged regions are shown in blue and red, respectively. **B.** Trx domains corresponding to the Proteins shown in Figure 1B. **C.** Sequence comparison of some selected vertebrate species and three *D. melanogaster* thioredoxins, (Tr-x2, TrxT and Dhd). RefSeq codes and species are indicated in Supplementary Table 1. The alignment was generated with Clustal Omega (EMBL-EBI) and the figure with BoxShade v3.21 (ExPASy). **D.** Extended version of the alignment of Dhd protein sequences shown in Figure 1C. The conservation level is indicated at the bottom of the alignment. Figure prepared with ESPript 3.0. **E.** Extended version of the alignment of TrxT protein sequences shown in Figure 1D. The conservation level is indicated at the bottom of the alignment. Figure prepared with ESPript 3.0. **F.** Comparison of divergent Dhd and Trx-2 sequences identified using Psi-Blast. Additional Lys and Arg residues present in Dhd but absents in TrxT are highlighted in blue.

**Supplementary Figure 2.**
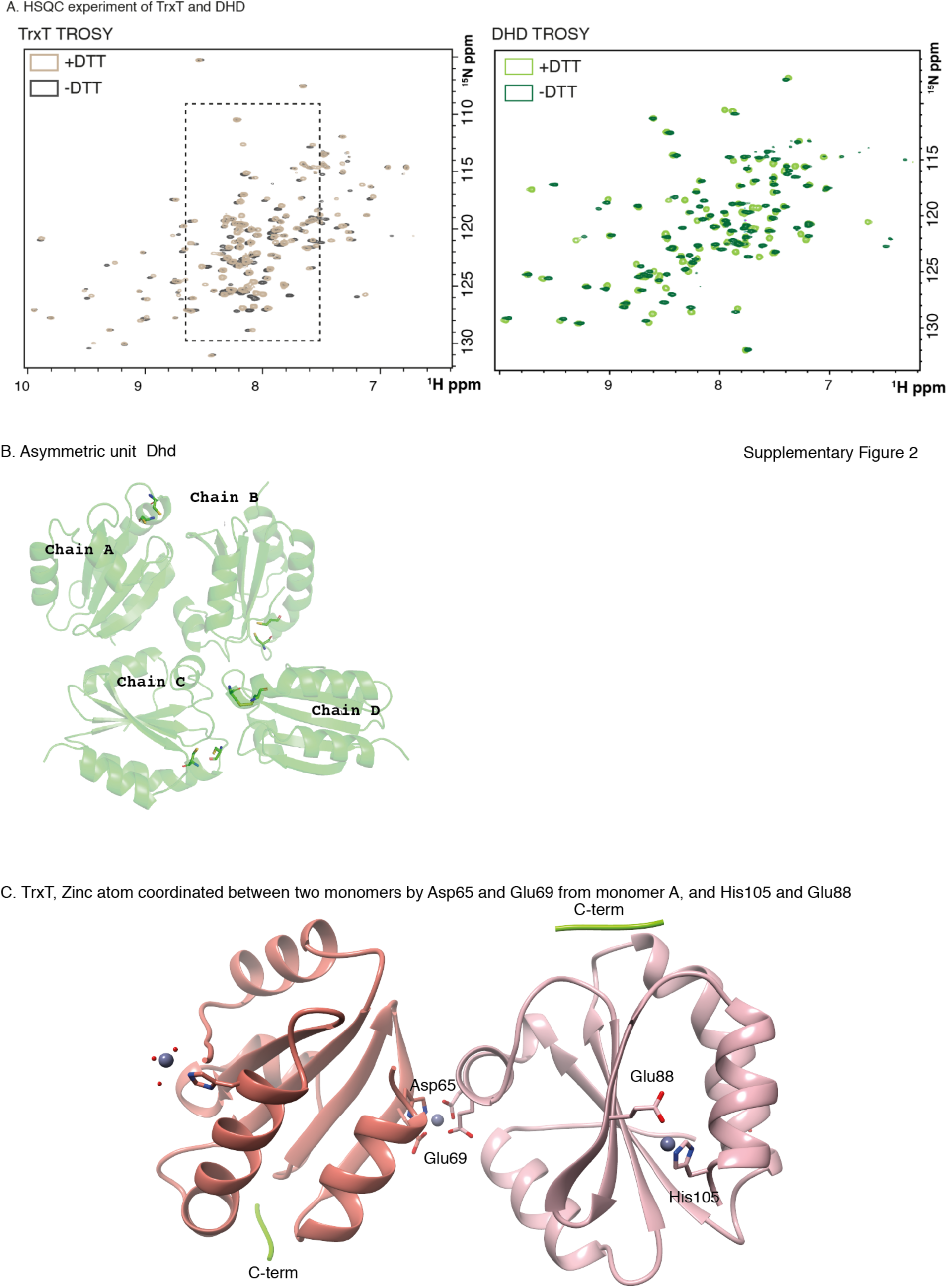
Structural characteristics of Dhd and TrxT by NMR and X-ray. **A.** 2D ^1^H, ^15^N Heteronuclear Single-Quantum Correlation HSQCHSQC experiments of TrxT and Dhd proteins, in the presence or absence of DTT. Chemical shift variations observed upon addition of DTT are interpreted as the results of modifications in the redox state of the proteins. The region containing the residues assigned to the C-terminal domain is indicated with a box. **B.** Cartoon representation of the asymmetric unit for the deadhead protein structure composed of four monomers. **C.** Cartoon representation of a symmetry-related dimer of the TrxT protein. Two monomers are engaged in a dimer interaction with symmetry-related neighbors through the coordination of a Zn atom. Residues involved in Zn interactions are labeled. The fragment of the C-terminal domain bound to Cys125 is shown in chartreuse. The rest of the C-terminal domain is not shown for simplicity.

**Supplementary Figure 3.**
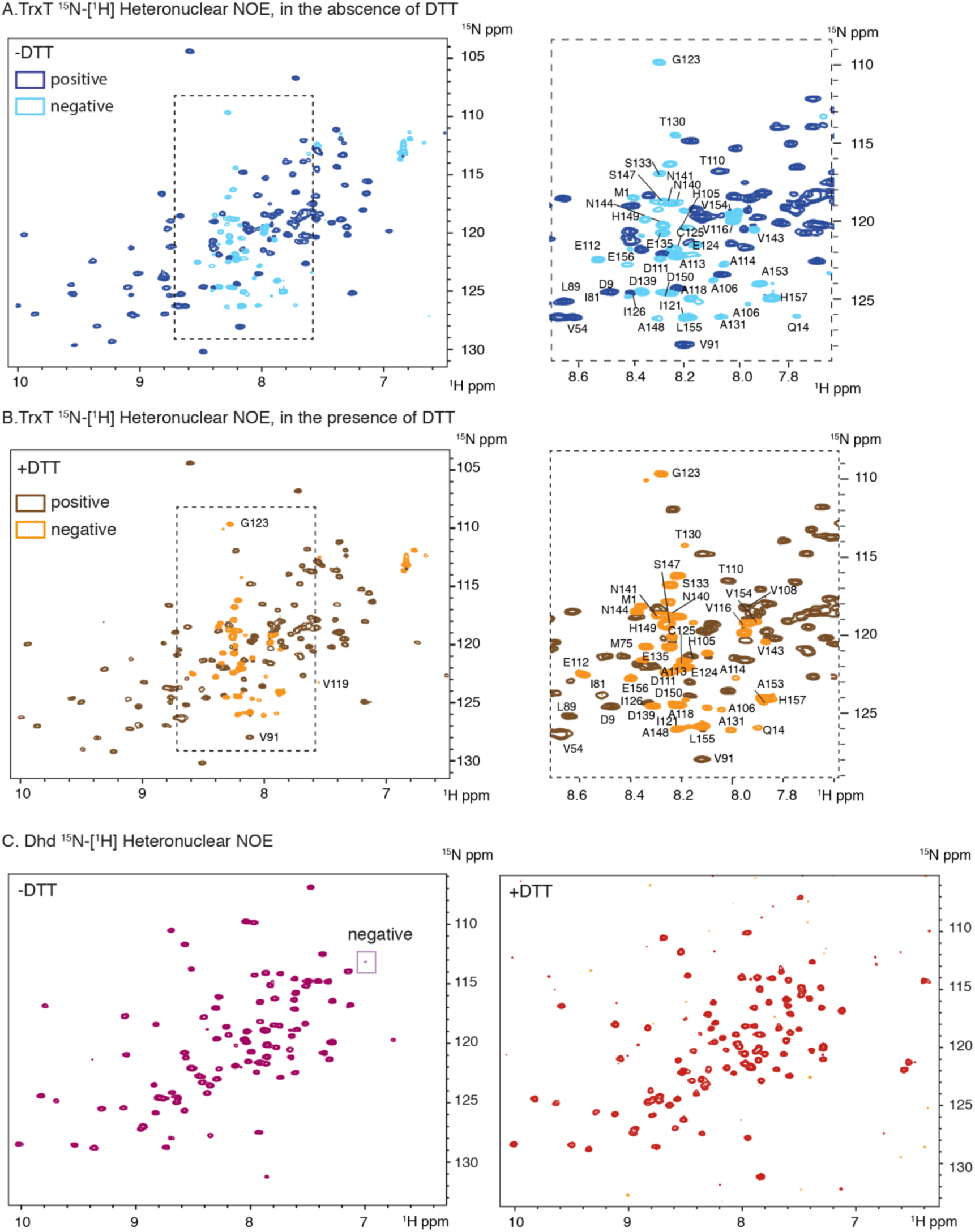
Flexible properties of the TrxT C-terminal domain. **A.** 2D ^15^N-[^1^H] heteronuclear NOEs (run as duplicates) indicating the positive and negative peaks of TrxT in the absence of DTT. Resonances corresponding to the C-terminal domain are labeled. A few corresponding to well-structured regions are also indicated for comparison. **B.** 2D ^15^N-[^1^H] heteronuclear NOEs (run as duplicates) in the presence of DTT. Some chemical shift variations are observed with respect to A, indicating the effect of DTT on the redox properties of the sample. In both cases, Cys125 displays negative NOEs, thereby indicating that its internal motion is not fully dependent on its redox state. **C.** Same experiments as those shown in A and B for Dhd. The fold is highly defined and negative peaks are not detected (with the exception of one side-chain).

**Supplementary Figure 4.**
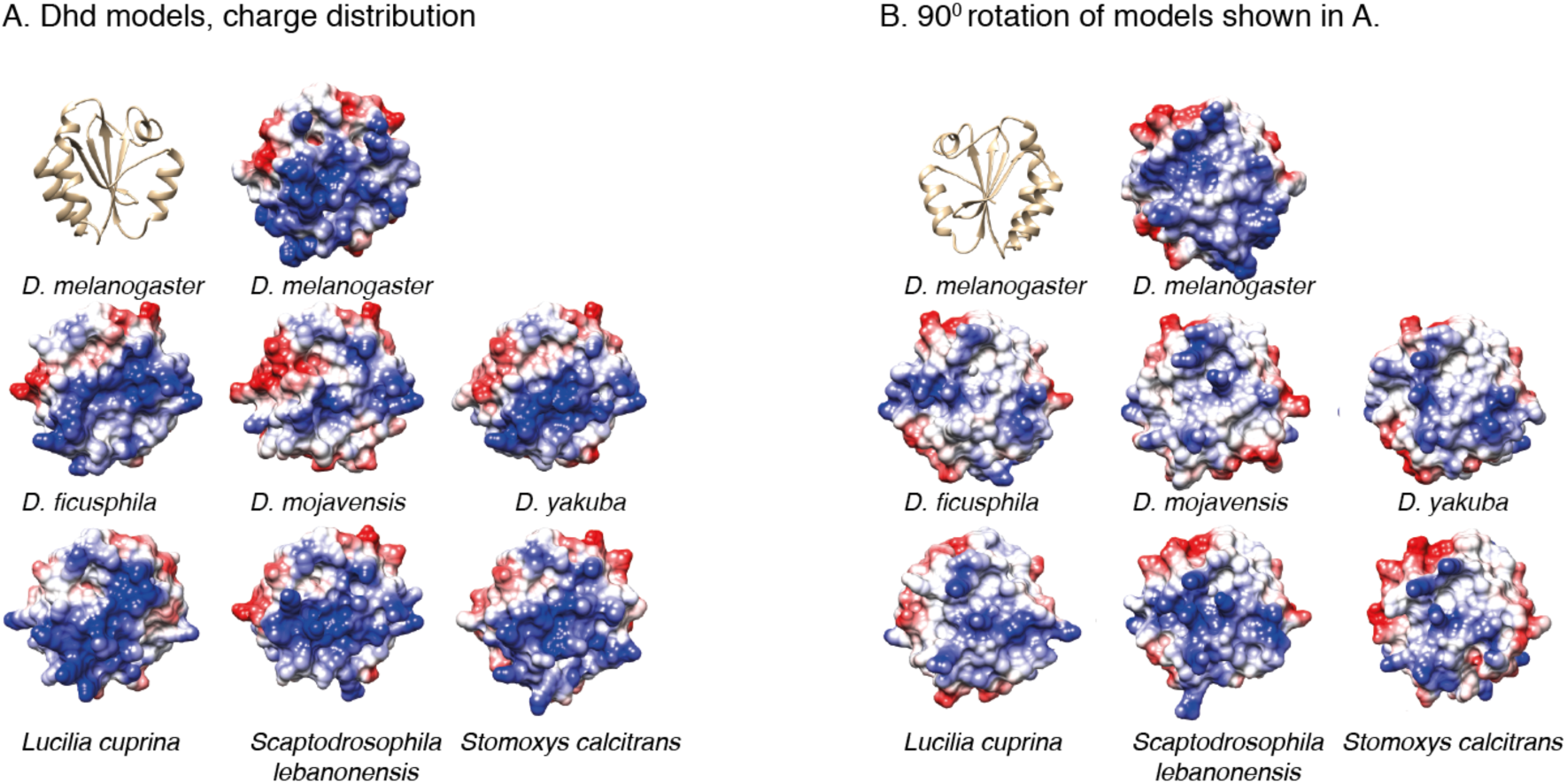
Charge distribution of Dhd models based on Dm Dhd structure. **A.** Structure and charge distribution of Dm Dhd (top) and of six additional Dhd sequences oriented like Dm Dhd. **B.** A 90-degree rotation of the surfaces displaying the other side of the molecule.

## Notes

### Competing Interest Statement

The authors have declared no competing interest.

## REFERENCES

1. Holmgren, A. (1985) Thioredoxin. Annu Rev Biochem, 54, 237–271.

2. Collet, J.F. and Messens, J. (2010) Structure, function, and mechanism of thioredoxin proteins. Antioxid Redox Signal, 13, 1205–1216.

3. Wilkinson, B. and Gilbert, H.F. (2004) Protein disulfide isomerase. Biochim Biophys Acta, 1699, 35–44.

4. Wahl, M.C., Irmler, A., Hecker, B., Schirmer, R.H. and Becker, K. (2004) Comparative Structural Analysis of Oxidized and Reduced Thioredoxin from Drosophila melanogaster. Journal of Molecular Biology, 345, 1119–1130.

5. Fomenko, D.E., Marino, S.M. and Gladyshev, V.N. (2008) Functional diversity of cysteine residues in proteins and unique features of catalytic redox-active cysteines in thiol oxidoreductases. Mol Cells, 26, 228–235.

6. Grogan, T.M., Fenoglio-Prieser, C., Zeheb, R., Bellamy, W., Frutiger, Y., Vela, E., Stemmerman, G., Macdonald, J., Richter, L., Gallegos, A. et al. (2000) Thioredoxin, a putative oncogene product, is overexpressed in gastric carcinoma and associated with increased proliferation and increased cell survival. Hum Pathol, 31, 475–481.

7. Smart, D.K., Ortiz, K.L., Mattson, D., Bradbury, C.M., Bisht, K.S., Sieck, L.K., Brechbiel, M.W. and Gius, D. (2004) Thioredoxin reductase as a potential molecular target for anticancer agents that induce oxidative stress. Cancer Res, 64, 6716–6724.

8. Bradshaw, T.D., Matthews, C.S., Cookson, J., Chew, E.H., Shah, M., Bailey, K., Monks, A., Harris, E., Westwell, A.D., Wells, G. et al. (2005) Elucidation of thioredoxin as a molecular target for antitumor quinols. Cancer Res, 65, 3911–3919.

9. Richardson, H.E.W.L.H.P.O. (2015) Screening for Anti-cancer Drugs in Drosophila. eLS.

10. Svensson, M.J., Chen, J.D., Pirrotta, V. and Larsson, J. (2003) The ThioredoxinT and deadhead gene pair encode testis- and ovary-specific thioredoxins in Drosophila melanogaster. Chromosoma, 112, 133–143.

11. Rossi, F., Molnar, C., Hashiyama, K., Heinen, J.P., Pamplona, J., Llamazares, S., Reina, J., Hashiyama, T., Rai, M., Pollarolo, G. et al. (2017) An in vivo genetic screen in Drosophila identifies the orthologue of human cancer/testis gene SPO11 among a network of targets to inhibit lethal(3)malignant brain tumour growth. Open Biology, 7, 170156.

12. Janic, A., Mendizabal, L., Llamazares, S., Rossell, D. and Gonzalez, C. (2010) Ectopic Expression of Germline Genes Drives Malignant Brain Tumor Growth in Drosophila. Science, 330, 1824–1827.

13. Ravi, D., Wiles, A.M., Bhavani, S., Ruan, J., Leder, P. and Bishop, A.J. (2009) A network of conserved damage survival pathways revealed by a genomic RNAi screen. PLoS Genet, 5, e1000527.

14. Emelyanov, A.V. and Fyodorov, D.V. (2016) Thioredoxin-dependent disulfide bond reduction is required for protamine eviction from sperm chromatin. Genes Dev, 30, 2651–2656.

15. Rathke, C., Baarends, W.M., Awe, S. and Renkawitz-Pohl, R. (2014) Chromatin dynamics during spermiogenesis. Biochim Biophys Acta., 155–168. .

16. Tirmarche, S., Kimura, S., Dubruille, R., Horard, B. and Loppin, B. (2016) Unlocking sperm chromatin at fertilization requires a dedicated egg thioredoxin in Drosophila. Nature Communications, 7, 13539.

17. Petrova, B., Liu, K., Tian, C., Kitaoka, M., Freinkman, E., Yang, J. and Orr-Weaver, T.L. (2018) Dynamic redox balance directs the oocyte-to-embryo transition via developmentally controlled reactive cysteine changes. Proc Natl Acad Sci U S A, 115, 7978–7986.

18. Petrova, B., Liu, K., Tian, C., Kitaoka, M., Freinkman, E., Yang, J. and Orr-Weaver, T.L. (2018) Dynamic redox balance directs the oocyte-to-embryo transition via developmentally controlled reactive cysteine changes. Proc Natl Acad Sci U S A, 115, E7978–E7986.

19. Torres-Campana, D., Kimura, S., Orsi, G.O., Horard, B., Bonoit, G. and Loppin, B. (2020) The Lid/KDM5 histone demethylase complex activates a critical effector of the oocyte-to-zygote transition. PLoS Genet, 16, e1008543.

20. Svensson, M.J., Stenberg, P. and Larsson, J. (2007) Organization and regulation of sex-specific thioredoxin encoding genes in the genus Drosophila. Dev Genes Evol, 217, 639–650.

21. Ubbink, M. (2009) The courtship of proteins: Understanding the encounter complex. FEBS Letters, 1060–1066.

22. Aragon, E., Wang, Q., Zou, Y., Morgani, S.M., Ruiz, L., Kaczmarska, Z., Su, J., Torner, C., Tian, L., Hu, J. et al. (2019) Structural basis for distinct roles of SMAD2 and SMAD3 in FOXH1 pioneer-directed TGF-beta signaling. Genes Dev, 33, 1506–1524.

23. Guca, E., Suñol, D., Ruiz, L., Konkol, A., Cordero, J., Torner, C., Aragon, E., Martin-Malpartida, P., Riera, T. and Macias, M.J. (2018) TGIF1 Homeodomain Interacts With Smad MH1 Domain and Represses TGF-β Signaling. Nucleic Acids Res, 46, 9220–9235.

24. Martin-Malpartida, P., Batet, M., Kaczmarska, Z., Freier, R., Gomes, T., Aragon, E., Zou, Y., Wang, Q., Xi, Q., Ruiz, L. et al. (2017) Structural basis for genome wide recognition of 5-bp GC motifs by SMAD transcription factors. Nat Commun, 8, 2070.

25. Marley, J., Lu, M. and Bracken, C. (2001) A method for efficient isotopic labeling of recombinant proteins. J Biomol NMR, 20, 71–75.

26. Sievers, F. and Higgins, D.G. (2018) Clustal Omega for making accurate alignments of many protein sequences. Protein Sci, 27, 135–145.

27. Robert, X. and Gouet, P. (2014) Deciphering key features in protein structures with the new ENDscript server. Nucleic Acids Res, 42, W320–324.

28. Drozdetskiy, A., Cole, C., Procter, J. and Barton, G.J. (2015) JPred4: a protein secondary structure prediction server. Nucleic Acids Research, 43, W389–W394.

29. Ishida, T. and Kinoshita, K. (2007) PrDOS: prediction of disordered protein regions from amino acid sequence. Nucleic Acids Research, 35, W460–W464.

30. Farrow, N.A., Muhandiram, R., Singer, A.U., Pascal, S.M., Kay, C.M., Gish, G., Shoelson, S.E., Pawson, T., Forman-Kay, J.D. and Kay, L.E. (1994) Backbone dynamics of a free and phosphopeptide-complexed Src homology 2 domain studied by 15N NMR relaxation. Biochemistry, 33, 5984–6003.

31. Kay, L.E., Torchia, D.A. and Bax, A. (1989) Backbone dynamics of proteins as studied by 15N inverse detected heteronuclear NMR spectroscopy: application to staphylococcal nuclease. Biochemistry, 28, 8972–8979.

32. Favier, A. and Brutscher, B. (2019) NMRlib: user*-*friendly pulse sequence tools for Bruker NMR spectrometers. ournal of Biomolecular NMR, 73, 199–211.

33. Bottomley, M.J., Macias, M.J., Liu, Z. and Sattler, M. (1999) A novel NMR experiment for the sequential assignment of proline residues and proline stretches in 13C/15N-labeled proteins. Journal of Biomolecular NMR volume 13, 381–385.

34. Delaglio, F., Grzesiek, S., Vuister, G.W., Zhu, G., Pfeifer, J. and Bax, A. (1995) NMRPipe : a multidimensional spectral processing system based on UNIX pipes. J Biomol NMR, 6, 277–293.

35. Vonrhein, C., Flensburg, C., Keller, P., Sharff, A., Smart, O., Paciorek, W., Womack, T. and Bricogne, G. (2011) Data processing and analysis with the autoPROC toolbox. Acta Crystallogr D Biol Crystallogr, 67, 293–302.

36. Tickle, I.J., Flensburg, C., Keller, P., Paciorek, W., Sharff, A., Vonrhein, C. and Bricogne, G. (2018) STARANISO. Global Phasing Ltd., Cambridge, UK.

37. McCoy, A.J. (2007) Solving structures of protein complexes by molecular replacement with Phaser. Acta Crystallogr D Biol Crystallogr, 63, 32–41.

38. McCoy, A.J., Grosse-Kunstleve, R.W., Adams, P.D., Winn, M.D., Storoni, L.C. and Read, R.J. (2007) Phaser crystallographic software. J Appl Crystallogr, 40, 658-674.

39. Murshudov, G.N., Vagin, A.A. and Dodson, E.J. (1997) Refinement of macromolecular structures by the maximum-likelihood method. Acta Crystallogr D Biol Crystallogr, 53, 240–255.

40. Liebschner, D., Afonine, P.V., Baker, M.L., Bunkoczi, G., Chen, V.B., Croll, T.I., Hintze, B., Hung, L.W., Jain, S., McCoy, A.J. et al. (2019) Macromolecular structure determination using X-rays, neutrons and electrons: recent developments in Phenix. Acta Crystallogr D Struct Biol, 75, 861–877.

41. Winn, M.D., Ballard, C.C., Cowtan, K.D., Dodson, E.J., Emsley, P., Evans, P.R., Keegan, R.M., Krissinel, E.B., Leslie, A.G., McCoy, A. et al. (2011) Overview of the CCP4 suite and current developments. Acta Crystallogr D Biol Crystallogr, 67, 235–242.

42. Emsley, P., Lohkamp, B., Scott, W.G. and Cowtan, K. (2010) Features and Development of Coot. Acta Crystallographica Section D - Biological Crystallography, 66, 486.

43. Pettersen, Goddard, Huang, Couch, Greenblatt, Meng and Ferrin. (2004) UCSF Chimera - a visualization system for exploratory research and analysis. J Comput Chem, 25, 1605–1612.

44. Madeira, F., Park, J.M., Lee, Y., Buso, N., Gur, T., Madhusoodanan, N., Basutkar, P., Tivey, A.R.N., Potter, S.C., Finn, R.D. et al. (2019) The EMBL-EBI search and sequence analysis tools APIs. Nucleic Acids Research, 47, W636–W641.

45. Feuerstein, S., Plevin, M.J., Willbold, D. and Brutscher, B. (2012) iHADAMAC: A complementary tool for sequential resonance assignment of globular and highly disordered proteins. Journal of Magnetic Resonance, 214, 329–334.

46. Sharma, D. and Rajarathnam, K. (2000) 13C NMR chemical shifts can predict disulfide bond formation. Journal of Biomolecular NMR, 18, 165–171.

47. Hwang, J., Suh, H.W., Jeon, Y.H., Hwang, E., Nguyen, L.T., Yeom, J., Lee, S.G., Lee, C., Kim, K.J., Kang, B.S. et al. (2014) The structural basis for the negative regulation of thioredoxin by thioredoxin-interacting protein. Nature Communications, 5, 2958.

48. Hwang, J., Suh, H.W., Jeon, Y.H., Hwang, E., Nguyen, L.T., Yeom, J., Lee, S.G., Lee, C., Kim, K.J., Kang, B.S. et al. (2014) The structural basis for the negative regulation of thioredoxin by thioredoxin-interacting protein. Nat Commun, 5, 2958.

49. Matthews, J.R., Wakasugi, N., Virelizier, J.L., Yodoi, J. and Hay, R.T. (1992) Thioredoxin regulates the DNA binding activity of NF-kappa B by reduction of a disulphide bond involving cysteine 62. Nucleic Acids Res., 20, 3821.

50. Rigobello, M.P. and Bindoli, A. (2010) Mitochondrial thioredoxin reductase purification, inhibitor studies, and role in cell signaling. Methods Enzymol, 474, 109–122.

51. Arner, E.S. and Holmgren, A. (2000) Physiological functions of thioredoxin and thioredoxin reductase. Eur J Biochem, 267, 6102–6109.

52. Powis, G., Mustacich, D. and Coon, A. (2000) The role of the redox protein thioredoxin in cell growth and cancer. Free Radic Biol Med, 29, 312–322.

53. Tamura, T. and Stadtman, T.C. (1996) A new selenoprotein from human lung adenocarcinoma cells: purification, properties, and thioredoxin reductase activity. Proc Natl Acad Sci U S A, 93, 1006–1011.

54. Eckenroth, B.E., Rould, M.A., Hondal, R.J. and Everse, S.J. (2007) Structural and biochemical studies reveal differences in the catalytic mechanisms of mammalian and Drosophila melanogaster thioredoxin reductases. Biochemistry, 46, 4694–4705.

55. Lagunas, A., Guerra-Castellano, A., Nin-Hill, A., Diaz-Montero, I., De la Rosa, M.A., Samitier, J., Rovira, C. and Gorostiza, P. (2018) Long distance electron transfer through the aqueous solution between redox partner proteins. Nature Communications 9, 5157.

56. Wang, C., Zhang, Q., Hang, T., Tao, Y., Ma, X., Wu, M., Zhang, X. and Zang, J. (2015) Structure of the JmjC domain-containing protein NO66 complexed with ribosomal protein Rpl8. Acta Crystallogr D Biol Crystallogr, 71, 1955–1964.

57. Du, Y., Zhang, H., Zhang, X., Lu, J. and Holmgren, A. (2013) Thioredoxin 1 is inactivated due to oxidation induced by peroxiredoxin under oxidative stress and reactivated by the glutaredoxin system. J Biol Chem, 288, 32241–32247.

58. Gonzalez, C. (2013) Drosophila melanogaster: a model and a tool to investigate malignancy and identify new therapeutics. Nat Rev Cancer, 13, 172–183.

59. Kozlowski, L.P. (2016) Isoelectric Point Calculator. Biology Direct, 11, 55.

